# Shared and unique brain network features predict cognition, personality and mental health in childhood

**DOI:** 10.1101/2020.06.24.168724

**Authors:** Jianzhong Chen, Angela Tam, Valeria Kebets, Csaba Orban, Leon Qi Rong Ooi, Scott Marek, Nico Dosenbach, Simon Eickhoff, Danilo Bzdok, Avram J Holmes, B.T. Thomas Yeo

**Affiliations:** Department of Electrical and Computer Engineering & N.1 Institute for Health, National University of Singapore, Singapore; Clinical Imaging Research Centre, National University of Singapore, Singapore; Centre for Sleep and Cognition, National University of Singapore, Singapore; NUS Graduate School for Integrative Sciences and Engineering, National University of Singapore, Singapore; Martinos Center for Biomedical Imaging, Massachusetts General Hospital, Charlestown, MA, USA; Department of Neurology, Washington University in St. Louis, St. Louis, MO, USA; Institute for Systems Neuroscience, Medical Faculty, Heinrich-Heine University Düsseldorf, Düsseldorf, Germany; Institute of Neuroscience and Medicine, Brain & Behaviours (INM-7), Research Center Jülich, Jülich, Germany; Department of Biomedical Engineering, Montreal Neurological Institute, McGill University, Montreal, Quebec, Canada; Mila - Quebec AI Institute, Montreal, Canada; Yale University, Departments of Psychology and Psychiatry, New Haven, CT, USA

## Abstract

The manner through which individual differences in brain network organization track population-level behavioral variability is a fundamental question in systems neuroscience. Recent work suggests that resting-state and task-state functional connectivity can predict specific traits at the individual level. However, the focus of most studies on single behavioral traits has come at the expense of capturing broader relationships across behaviors. Here, we utilized a large-scale dataset of 1858 typically developing children to estimate whole-brain functional network organization that is predictive of individual differences in cognition, impulsivity-related personality, and mental health during rest and task states. Predictive network features were distinct across the broad behavioral domains: cognition, personality and mental health. On the other hand, traits within each behavioral domain were predicted by highly similar network features. This is surprising given decades of research emphasizing that distinct brain networks support different mental processes. Although tasks are known to modulate the functional connectome, we found that predictive network features were similar between resting and task states. Overall, our findings reveal shared brain network features that account for individual variation within broad domains of behavior in childhood, yet are unique to different behavioral domains.

## Introduction

A central question in systems neuroscience is how brain network architecture supports the wide repertoire of human behavior across the lifespan. Childhood is a period of rapid neural development and behavioral changes across cognition, personality, and mental health (Steinberg 2005, Casey *et al.* 2008, Paus *et al.* 2008). Consequently, there is particular interest in understanding the nature of brain-behavior relationships instantiated early in the lifespan (Spear 2013, Larsen and Luna 2018). Here, we utilized a large-scale dataset of typically developing 9- to 10-year-old children (Volkow *et al.* 2018) to quantitatively characterize functional network organization that supports individual-level prediction of cognition, impulsivity-related personality, and mental health across resting and task states.

Whole-brain connectome-wide neurodevelopmental studies have found associations between resting-state functional network organization and behavioral traits (Satterthwaite *et al.* 2015, Karcher *et al.* 2019, Marek *et al.* 2019, Pornpattananangkul *et al.* 2019). However, clinical decisions are made at the individual level (Milham *et al.* 2017, Bzdok and Meyer-Lindenberg 2018). As such, there is an increasing shift from associational analyses to individual-level prediction (Dosenbach *et al.* 2010, Finn *et al.* 2015, Hsu *et al.* 2018, Nostro *et al.* 2018, Kong *et al.* 2019). Using machine learning algorithms, we can exploit inter-individual heterogeneity in functional connectomes to make predictions about a single person’s behavior (Finn *et al.* 2015). Consequently, neurodevelopmental prediction studies have used resting-state functional connectivity (FC) to predict individual differences in cognition (Evans *et al.* 2015, Sripada *et al.* 2019, Cui *et al.* 2020), impulsivity (Shannon *et al.* 2011) and autism symptoms (Uddin *et al.* 2013, Lake *et al.* 2019).

Recent studies have further suggested that task-state FC yields better prediction of cognition over resting-FC (Rosenberg *et al.* 2016, Greene *et al.* 2018, Jiang *et al.* 2019), with additional performance improvements from combining task-FC and resting-FC (Elliott *et al.* 2019, Gao *et al.* 2019). The improvements suggest that functional connections predictive of individual-level cognition (i.e., predictive network features) might differ between rest and task states. However, other studies have shown that the brain functional network architecture is broadly similar during rest and task (Smith *et al.* 2009, Cole, Bassett, *et al.* 2014, Krienen *et al.* 2014). Indeed, while task contexts reliably modulate functional network organization (Schultz and Cole 2016, Shine *et al.* 2016, Salehi *et al.* 2019), task modulation of the functional connectome within individuals is much smaller than differences between individuals (Gratton *et al.* 2018). Therefore, it remains unclear whether predictive network features differ across brain states. This is a central question we seek to address in this study.

Furthermore, most previous connectome-based prediction studies have focused on specific behavioral traits (Rosenberg *et al.* 2016, Greene *et al.* 2018, Nostro *et al.* 2018, Wang *et al.* 2018, Jiang *et al.* 2019, Lake *et al.* 2019, Sripada *et al.* 2019, Cui *et al.* 2020). Yet, the human brain has evolved to execute a diverse range of behaviors, so focusing on single behavioral traits might miss the forest for the trees (Holmes and Patrick 2018). More specifically, it remains unclear whether predictive network features are similar or different across behavioral measures. For example, specialized brain networks support distinct cognitive processes, such as attention, language or attention (Corbetta and Shulman, 2002; Fedorenko and Thompson-Schill 2014; DiNicola et al., 2020). Thus, one might expect distinct network features to support prediction of different cognitive traits. On the other hand, many studies have also emphasized information integration across specialized brain networks (van den Heuvel and Sporns 2011, Cole *et al.* 2013, Bertolero *et al.* 2018). Consequently, one might also expect a common set of predictive network features that explain individual differences in cognition. To systematically revisit the two possible scenarios, we considered the prediction of a large number of behavioral measures. This population neuroscience re-assessment allowed us to estimate the degree of overlap in predictive network features across different behavioral domains (cognition, personality, mental health), as well as across phenotypes within the same behavioral domain.

In the present study, we utilized the Adolescent Brain Cognitive Development (ABCD) study, a unique dataset with a large sample of children and a diverse set of behavioral measures (Volkow *et al.* 2018). We used resting-FC and task-FC to predict a wide range of cognitive, impulsivity-related personality, and mental health measures. We also investigated whether combining resting-FC and task-FC can improve behavioral prediction. Most importantly, we explored the existence of shared and unique predictive network features within and across behavioral domains, as well as across brain (resting and task) states.

## Results

We used resting-fMRI and task-fMRI from 11875 children (ABCD 2.0.1 release). There were three tasks: monetary incentive delay (MID), stop signal task (SST) and N-Back. We also considered all available dimensional neurocognitive (Luciana *et al.* 2018) and mental health (Barch *et al.* 2018) assessments, yielding 16 cognitive, 11 (impulsivity-related) personality and 9 mental health measures. After strict preprocessing quality control (QC) and considering only participants with complete resting-fMRI, task-fMRI and behavioral data, our main analyses utilized data from 1858 unrelated children (Figure 1A).

**Figure 1.**
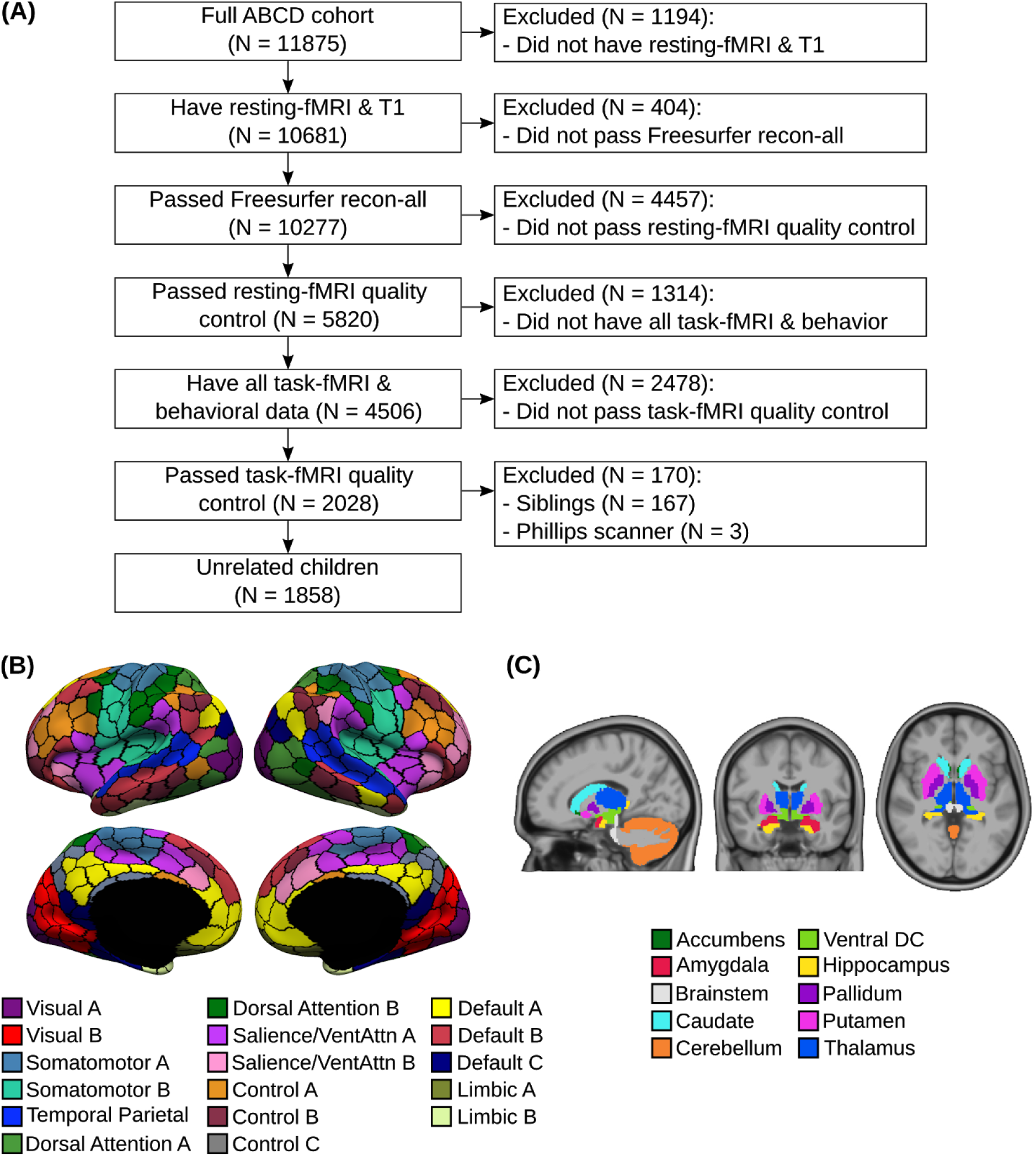
Overview of preprocessing workflow. (A) Flowchart illustrating inclusion/exclusion criteria. (B) Cortical parcellation of 400 regions (Schaefer *et al.* 2018). Parcel colors are assigned according to 17 large-scale networks (Yeo *et al.* 2011). (C) Nineteen subcortical regions (Fischl *et al.* 2002). Panels B and C were reproduced from Orban and colleagues (2020).

### Task-FC outperforms resting-FC for predicting cognition, but not personality or mental health

We computed FC (Pearson’s correlations) among the average time courses of 400 cortical (Schaefer *et al.* 2018) and 19 subcortical (Fischl *et al.* 2002) regions (Figures 1B & 1C), yielding a 419 × 419 FC matrix for each brain state (rest, MID, SST, N-back). We used kernel regression to predict each behavioral measure based on resting-FC, MID-FC, SST-FC and N-back-FC separately. We have previously demonstrated that kernel regression is a powerful approach for resting-FC behavioral prediction (He *et al.* 2020). The idea behind kernel regression is that subjects with more similar FC matrices would exhibit more similar behavior.

To evaluate the kernel regression performance, we utilized an inner-loop (nested) cross-validation procedure in which participants were repeatedly divided in training and test sets. The regression model was fitted on the training set and used to predict behavior in the test set. Care was taken so that participants from the same site were not split between training and test sets. This cross-validation procedure was repeated 120 times to ensure stability (Varoquaux *et al.* 2017). See Methods for more details.

Figure 2A shows the prediction performance averaged within each behavioral domain. Each behavioral domain was predicted better than chance (FDR q < 0.05) with p < 0.0005 across all brain states for cognition, (impulsivity-related) personality and mental health respectively.

**Figure 2.**
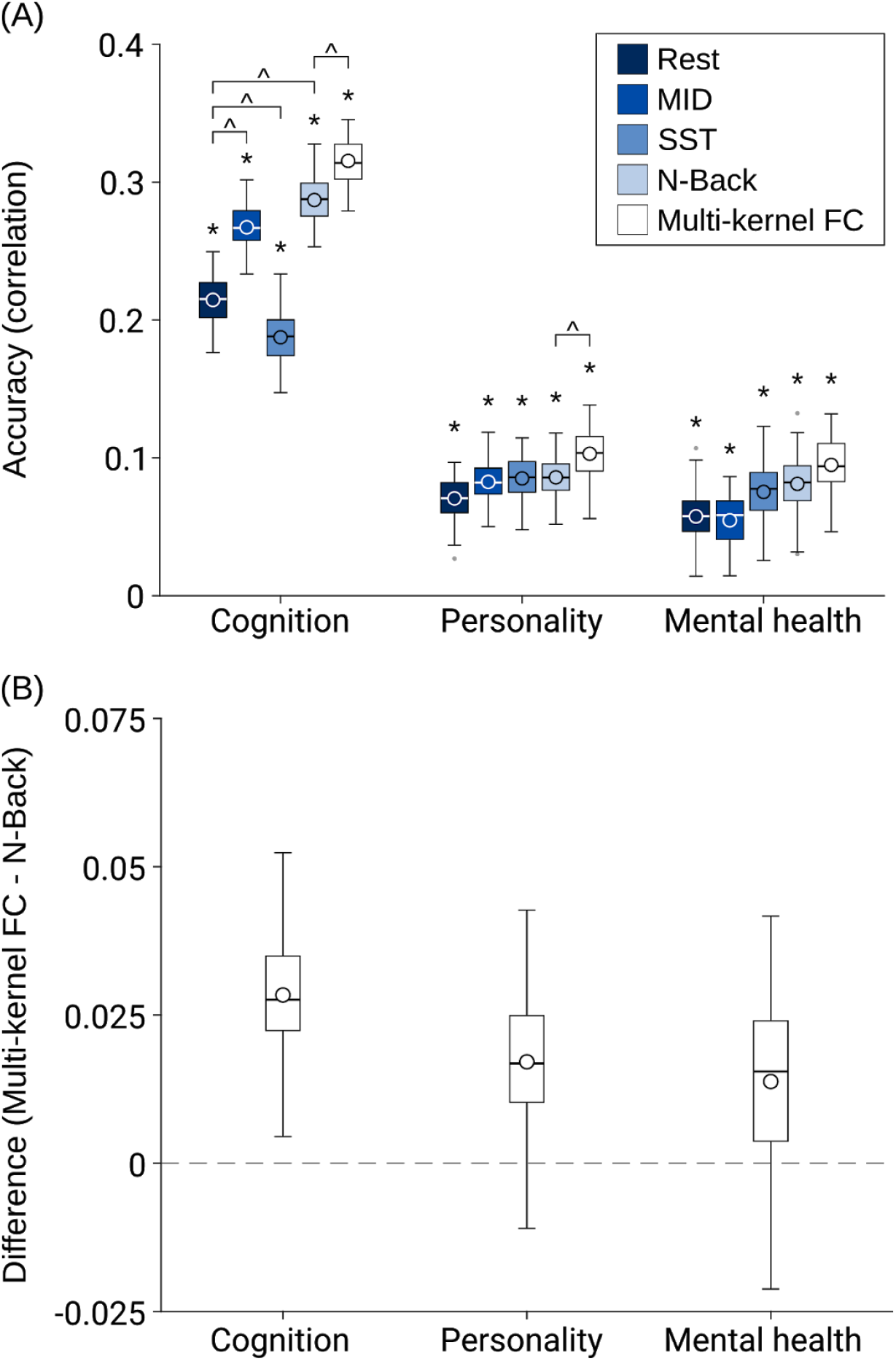
(A) Cross-validated prediction performance (Pearson’s correlation between observed and predicted values) using kernel ridge regression for resting-state and task-states (MID, SST, N-Back). Multi-kernel FC utilized FC from all 4 brain states for prediction. * denotes above chance prediction after correction for multiple comparisons (FDR q < 0.05). ^ denotes significantly different comparison after correction for multiple comparisons (FDR q < 0.05). The boxplots show the average accuracy across 120 replications. Task-FC appeared to only improve prediction performance for cognition, but not (impulsivity-related) personality or mental health. Multi-kernel FC improved prediction performance for cognition and personality, but not mental health. Similar conclusions were obtained using coefficient of determination (COD) instead of Pearson’s correlation as a measure of prediction performance (Figure S1). MID: monetary incentive delay; SST: stop signal task. (B) The average difference in accuracy (Pearson’s correlation between observed and predicted values) between the Multi-kernel FC and N-back models across 120 replications.

Consistent with previous studies (Greene *et al.* 2018), we found that MID-FC and N-back-FC outperformed resting-FC (p = 7.08e-08 and p = 4.85e-09 respectively) in predicting cognition. However, SST-FC had worse performance than resting-FC (p = 0.0082). In the case of personality and mental health, there was no statistical difference between resting-FC and any task state. Thus, task-FC appeared to improve prediction performance for cognition, but not personality or mental health.

### Combining task-FC and resting-FC improves prediction of cognition and personality, but not mental health

Previous studies have suggested that combining task-FC and resting-FC can improve prediction of fluid intelligence (Elliott *et al.* 2019, Gao *et al.* 2019) and reading comprehension (Jiang *et al.* 2019). We extended the previous studies by performing multi-kernel ridge regression using resting-FC, MID-FC, SST-FC and N-back-FC jointly to predict a broader range of cognitive measures as well as non-cognitive (personality and mental health) measures.

Figure 2 shows the multi-kernel prediction performance averaged within each behavioral domain. Since N-back performed the best among the single-kernel regression for all behavioral domains (Figure 2A), we compared multi-kernel FC with N-back-FC (Figure 2B). We found that multi-kernel FC performed better than N-back-FC for cognitive (p = 5.27e-06) and personality (p = 0.02), but not mental health (p = 0.12).

Figure 3 shows the prediction performance of multi-kernel FC for all individual behaviors. As can be seen, the prediction performance varies widely across behavioral measures. All 16 cognitive and 9 personality measures were significantly predicted better than chance, while 7 out of 11 mental health measures were significantly predicted. On average, across behavioral measures that were predicted better than chance, the correlation between observed and predicted values for cognition was 0.316 ± 0.126 (mean ± std), personality was 0.103 ± 0.044 and mental health was 0.120 ± 0.064.

**Figure 3.**
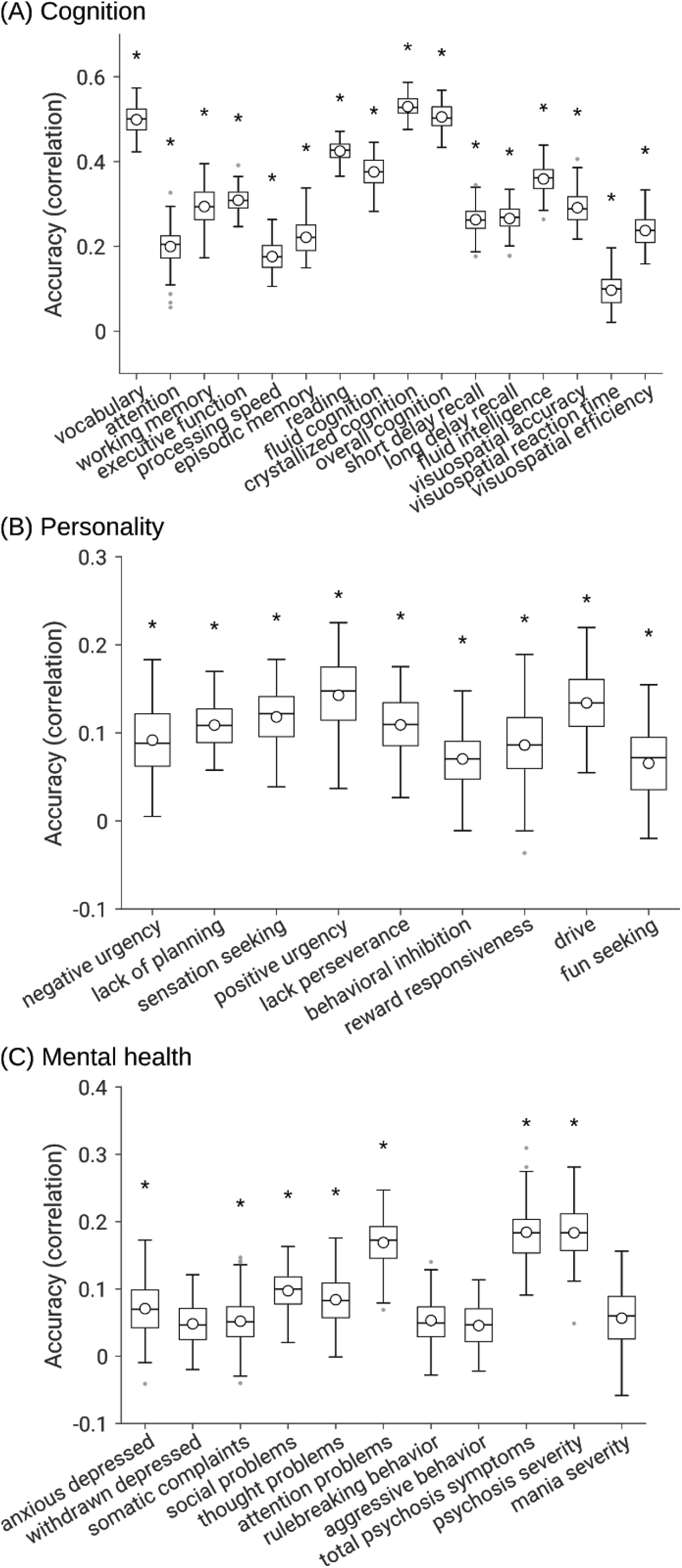
Cross-validated prediction performance (Pearson’s correlation between observed and predicted values) using multi-kernel ridge regression by exploiting resting-FC, MID-FC, SST-FC and N-back-FC jointly. (A) Cognitive measures. (B) (Impulsivity-related) Personality measures. (C) Mental health measures. * denotes above chance prediction after correcting for multiple comparisons (FDR q < 0.05). The boxplots show the average accuracy across 120 replications. Note the different scales across the three panels. The same set of behavioral measures were predicted better than chance when using coefficient of determination (COD) instead of Pearson’s correlation as a measure of prediction performance (Figure S2).

Thus, prediction performance was better for cognition than personality or mental health. For example, the best predicted cognitive measure was crystallized cognition with an accuracy of r = 0.530, while the best predicted personality measure was positive urgency with an accuracy of 0.143 and the best predicted mental health measure was total psychosis symptoms with an accuracy of 0.184. Henceforth, we will focus on the 32 behavioral measures that were significantly predicted by multi-kernel FC.

### Predictive brain network features cluster together within behavioral domains across all brain states

Most previous studies have focused on predicting a small number of behavioral measures. By considering a large number of behavioral measures across multiple behavioral domains, we were able to explore the question of whether predictive brain network features were shared or unique across behavioral measures. The multi-kernel regression models were inverted (Haufe *et al.* 2014), yielding a 419 × 419 predictive-feature matrix for each brain state (rest, MID, SST, N-back) and each behavioral measure. Haufe's inversion approach yields a positive (or negative) predictive-feature value for an edge, indicating that higher FC for the edge was associated with predicting greater (or lower) behavioral values. Figure 4 shows the predictive-feature matrices for positive urgency and negative urgency across all brain states. All predictive-feature matrices can be found in Figures S3 to S6.

**Figure 4.**
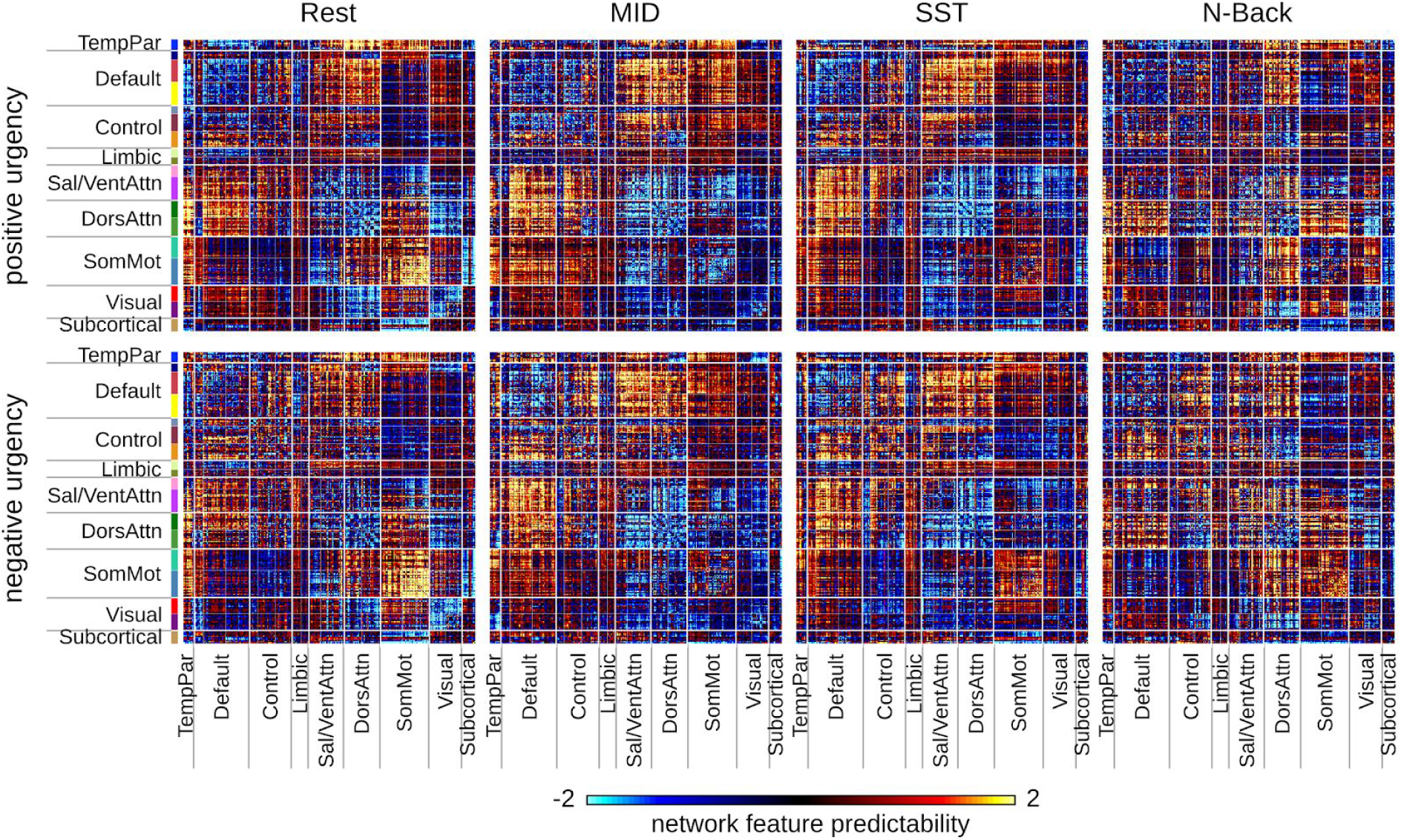
Predictive network features for positive urgency and negative urgency across all brain states. Haufe’s approach was utilized to invert the kernel regression models (Haufe *et al.* 2014), which allowed us to interpret which features were important for predicting a particular behavior. A positive (or negative) predictive-feature value indicates that higher FC was associated with predicting greater (or lower) behavioral values. As can be seen, the predictive features were similar between positive urgency and negative urgency across all brain states (although there were also some differences), motivating further analyses (Figures 5 and 6). Predictive-feature matrices for all behavioral measures can be found in Figures S3 to S6. For visualization, the values within each matrix were divided by their standard deviations. MID: monetary incentive delay; SST: stop signal task.

This inversion process is critical to interpreting supervised prediction models. Most previous studies have either interpreted the model weights or selected features, which leads to less interpretable results that are sensitive to the choice of regression models (Haufe et al., 2014). As will be shown in additional control analyses, we showed that the predictive features were highly robust across regression models, underlining the importance of this inversion process.

As can be seen in Figure 4, the predictive features were very similar between positive urgency and negative urgency for each brain state. The predictive features were also similar across brain states, but to a lower extent than the between-behavior similarity. To more quantitatively explore these phenomena, we first investigated whether predictive network features were similar across behavioral measures. Predictive-feature matrices for each behavioral measure were concatenated across brain states and correlated between behaviors, yielding a 32 × 32 matrix shown in Figure 5A. Here, the behavioral measures are ordered based on ABCD’s classification of these measures into cognition, personality and mental health behavioral domains, so we referred to this ordering as “hypothesis-driven”. If a pair of behavioral measures exhibited a high value (green) in the matrix (Figure 5A), then this indicates that the two behavioral measures are predicted by highly similar network features. As can be seen, the predictive-feature matrices were much more similar within each behavioral domain than across behavioral domains (Figure 5A).

**Figure 5.**
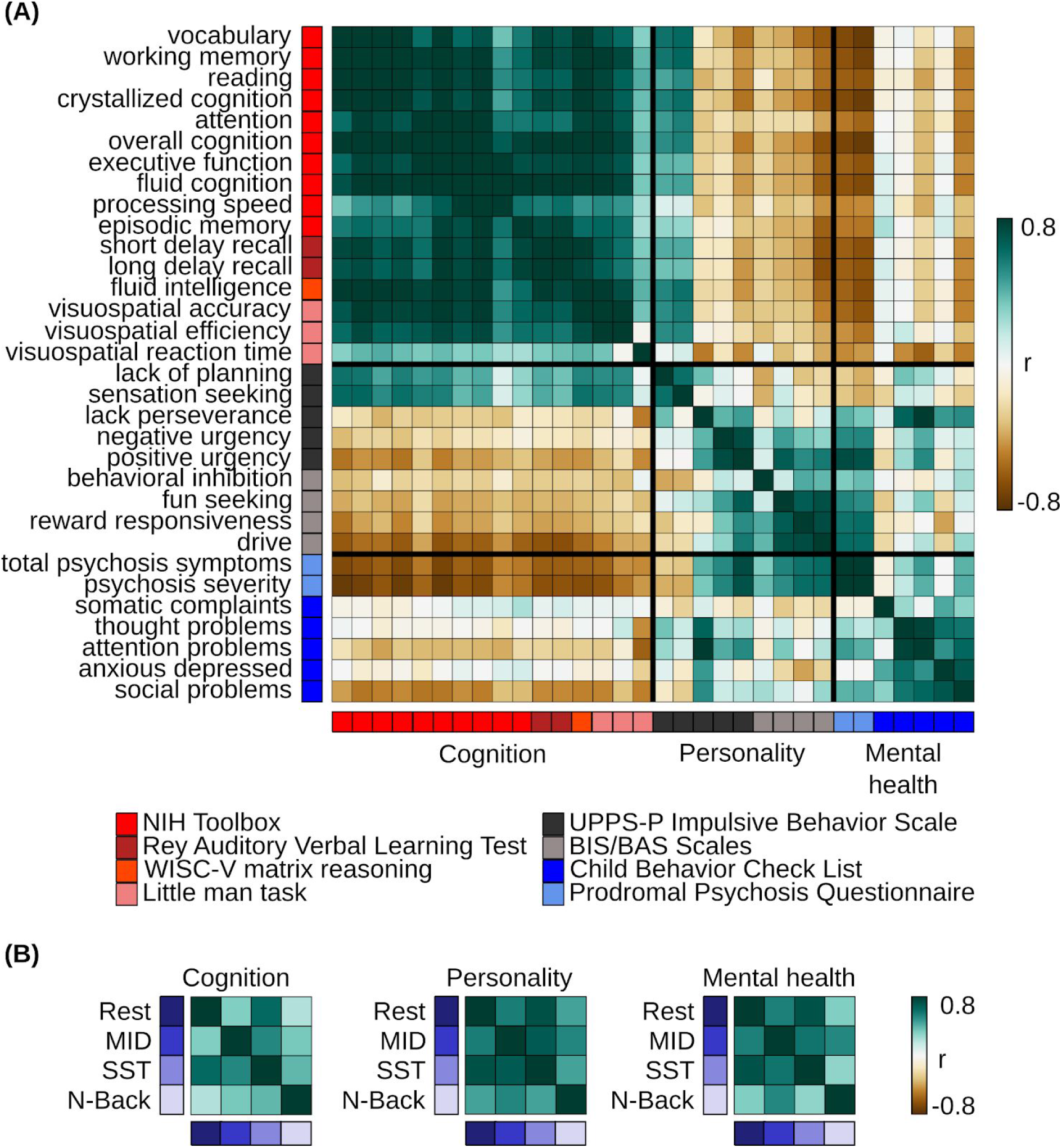
Predictive network features are similar within hypothesis-driven behavioral domains and across brain states. (A) Correlations of predictive-feature matrices (Figure 4) across behavioral measures. The predictive-feature matrices were concatenated across brain states and correlated across behavioral measures. If a pair of behavioral measures exhibited a high value (green), then this indicates that the two behavioral measures are predicted by highly similar network features. (B) Correlations of predictive-feature matrices across brain states. Predictive-feature matrices were averaged within each behavioral domain and correlated across brain states. The behavioral measures were ordered and categorized based on ABCD’s classification of these measures into cognition, personality and mental health behavioral domains, so we referred to this ordering as “hypothesis-driven”. Figure S10 shows the analogue of this figure, but without collapsing across either dimension of brain state or behavior. MID: monetary incentive delay; SST: stop signal task.

Instead of ordering the behavioral measures in a hypothesis-driven fashion (Figure 5A), we also re-ordered the behavioral measures by hierarchical clustering of the predictive-feature matrices (Figure 6A). The hierarchical clustering yielded three data-driven behavioral clusters (Figure 6A) that were highly similar to the hypothesis-driven behavioral domains (Figure 5A). We again see that the predictive-feature matrices were much more similar within each data-driven behavioral domain than across domains

**Figure 6.**
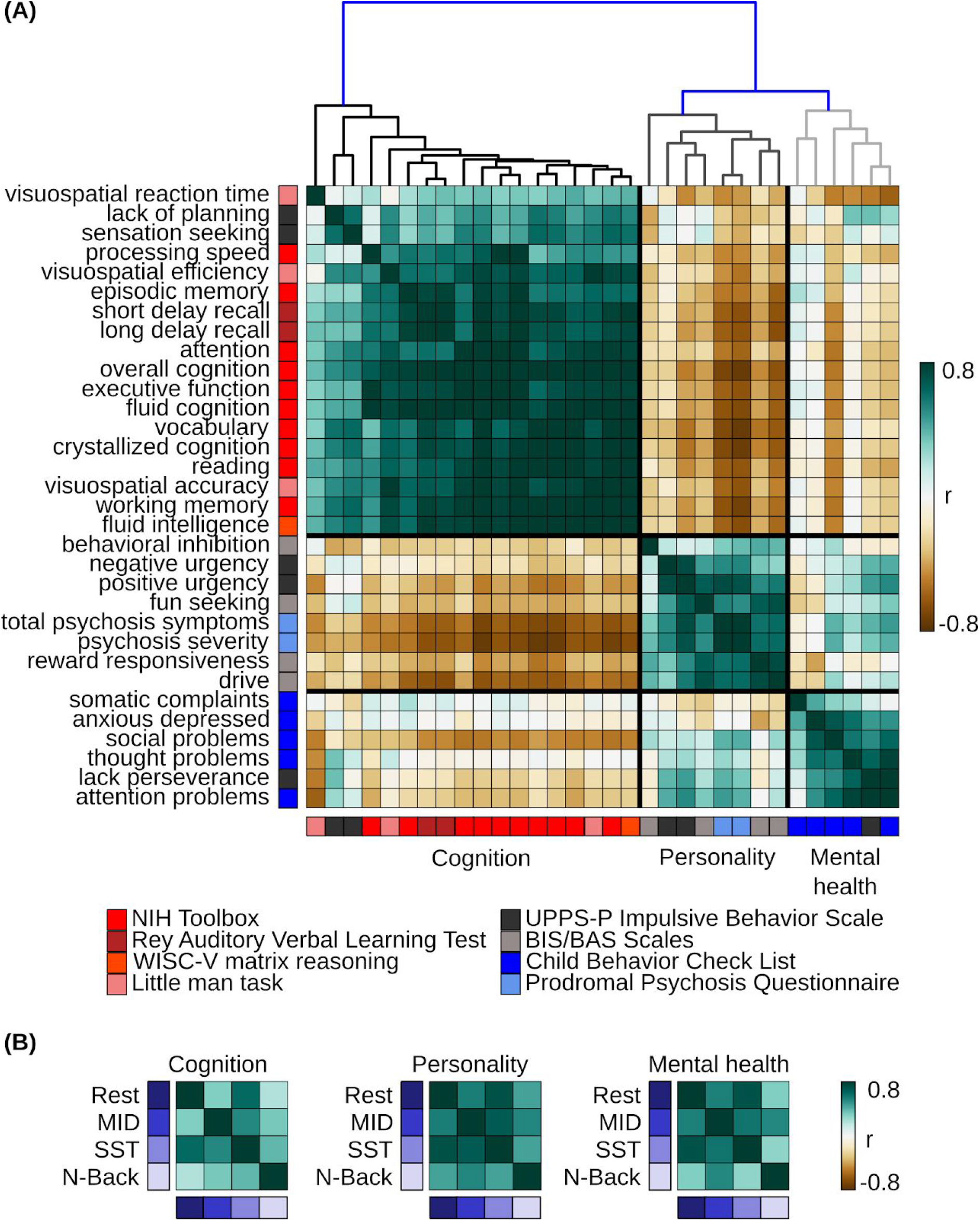
Predictive network features are similar within data-driven behavioral domains and across brain states. Both panels (A) and (B) are the same as Figure 5, except that behavioral measures are ordered and categorized based on the data-driven clusters of cognition, personality and mental health. These data-driven clusters were obtained by hierarchical clustering of the predictive-feature matrices (Figure 4) as indicated by the dendrogram in panel A. Clustering was performed using hierarchical agglomerative average linkage (UPGMA) clustering as implemented in scipy 1.2.1 (Virtanen *et al.* 2020). Figure S11 shows the analogue of this figure, but without collapsing across either dimension of brain state or behavior. MID: monetary incentive delay; SST: stop signal task.1

We then tested whether predictive network features were similar across brain states. Since predictive-feature matrices were similar within each behavioral domain (Figure 5A), we averaged the predictive-feature matrices across behaviors, yielding a predictive-feature matrix for each behavioral domain and each brain state (Figure S7). The 12 predictive-feature matrices were then correlated across behavioral domains and brain states. The predictive-feature matrices were similar across brain states within each behavioral domain (especially in the case of personality and mental health) (Figure 5B). Performing the same analyses using the data-driven behavioral clusters (Figures 6 & S8) yielded similar results.

Overall, these results suggest that predictive network features were more similar within behavioral domains (cognition, personality, mental health) than across behavioral domains. Furthermore predictive network features were similar across brain states. Critically, the similarity in predictive network features cannot be completely explained by similarity among the actual behavioral measures themselves (Figure S9). For example, “lacking of planning” and “sensation seeking” shared predictive features with cognitive measures (Figure 6A), although the behavioral measures themselves were more correlated with other mental health and personality measures (Figure S9). As another example, the average correlation of predictive network features across cognitive measures was 0.68 ± 0.19 (mean ± std), while the correlation among the raw cognitive scores was 0.29 ± 0.22.

### Distinct brain network features support the prediction of cognition, personality, and mental health

Having established that predictive network features were similar within behavioral domains and across brain states, we investigated the topography of predictive network features that were shared across states within each behavioral domain. Predictive-feature matrices were averaged within each hypothesis-driven behavioral domain, yielding 12 predictive-feature matrices (one for each behavioral domain and each brain state; Figure S7). To limit the number of multiple comparisons, permutation tests were performed for each within-network and between-network block by averaging predictive-feature values within and between 18 networks (FDR q < 0.05; Figure S12).

To examine predictive features common across brain states, we averaged the predictive-feature matrices across all brain states, considering only network blocks that were significant and exhibited the same directionality across states (Figure 7A). This conjunction thus highlights predictive network features that are shared across brain states and across behavioral measures within a behavioral domain. Figure 7B illustrates the connectivity strength obtained from averaging within each significant block. Figures 7C and 7D illustrate the predictability of each cortical region obtained by summing the rows of Figure 7A for positive and negative predictive-feature values separately (see subcortical regions in Figure S13A). As can be seen (Figures 7 & S13A) and consistent with the previous section (Figures 5 & 6), the patterns of predictive network features were distinct across the three behavioral domains.

**Figure 7.**
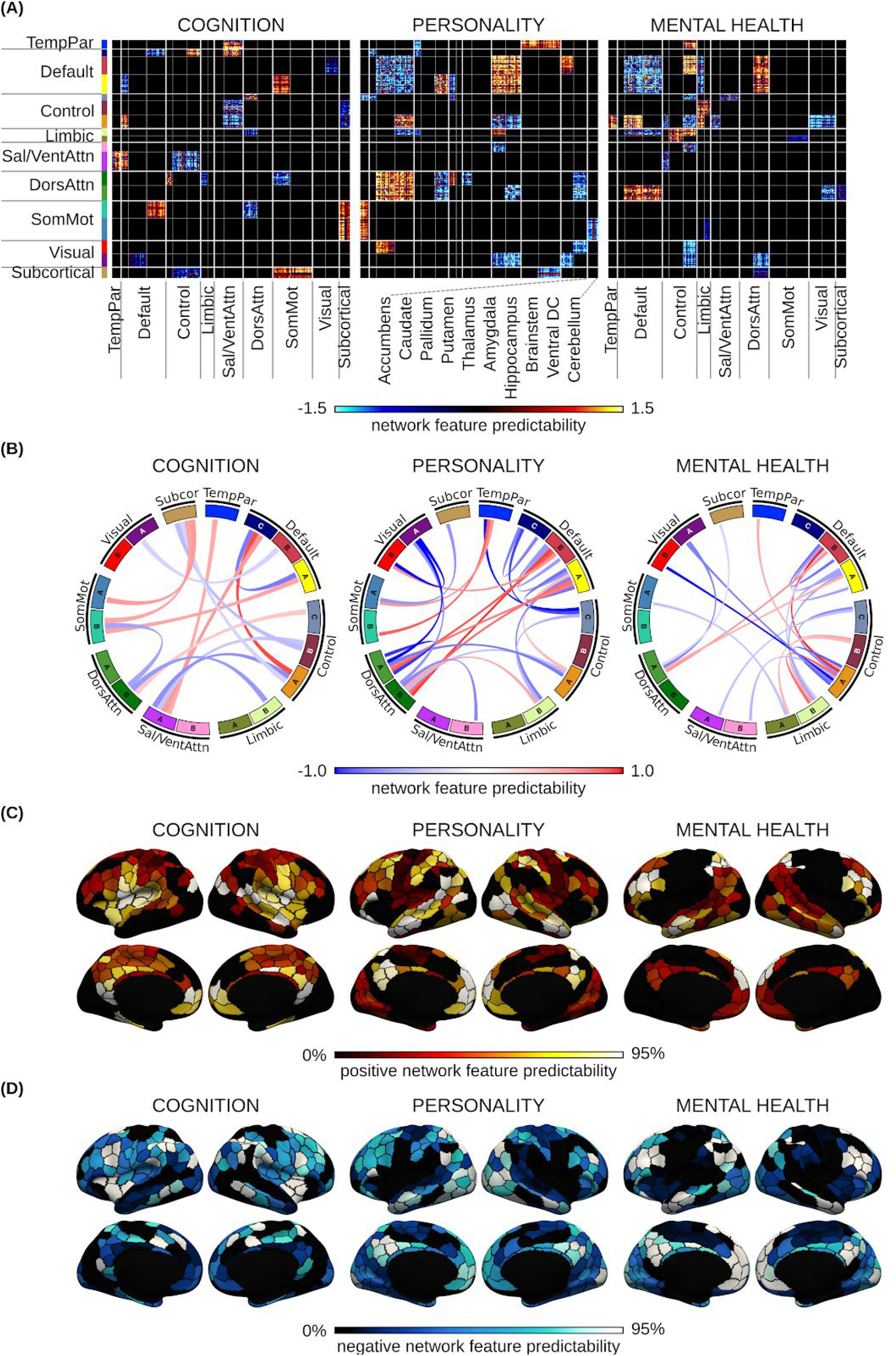
Brain network features that support individual-level prediction of cognition, personality and mental health. (A) Predictive-feature matrices averaged across brain states, considering only within-network and between-network blocks that were significant across all four brain states (Rest, MID, SST, N-Back). (B) Predictive network connections obtained by averaging the matrices in panel (A) within each between-network and within-network block. (C) Positive predictive features obtained by summing positive predictive-feature values across the rows of panel (A). A higher value for a brain region indicates that stronger connectivity yielded a higher prediction for the behavioral measure. (D) Negative predictive features obtained by summing negative predictive-feature values across the rows of panel (A). A higher value for a brain region indicates that weaker connectivity yielded a greater prediction for the behavioral measure. See Figure S13A for the subcortical maps. For visualization, the values within each matrix in panel A were divided by their standard deviations. The current figure utilized hypothesis-driven behavioral domains. Conclusions were highly similar using data-driven behavioral clusters (Figure S15).

Cognitive performance of individual participants was predicted by a distributed set of large-scale network features (Figures 7A & 7B) with somatomotor and salience networks being particularly prominent (Figures 7C & 7D). For example, lower connectivity of somatomotor network B with subcortical and default network A regions was predictive of higher cognitive scores (i.e., better cognition). As another example, greater connectivity between salience/ventral attention network A and default network C, as well as lower connectivity between salience/ventral attention network A and control networks were predictive of better cognition.

Personality measures of individual participants were predicted by a distributed set of large-scale network features (Figures 7A & 7B) with default and dorsal attentional networks being particularly prominent (Figures 7C & 7D). For example, greater connectivity between default networks A/B and dorsal attention networks A/B were predictive of greater personality scores (i.e., greater impulsivity and sensitivity to reward/punishment). On the other hand, lower connectivity within default networks A/B were predictive of greater impulsivity-related traits.

Mental health of individual participants was predicted by a distributed set of large-scale network features (Figures 7A & 7B) with default and frontoparietal control networks being particularly prominent (Figures 7C & 7D). For example, greater connectivity between default networks A/B and dorsal attention networks A/B were predictive of larger mental health scores (i.e., worse mental health). On the other hand, lower connectivity within default networks A/B were predictive of worse mental health.

As a control analysis, we utilized the previously derived data-driven clusters of cognition, personality and mental health (Figure 6) to perform the same analyses, yielding highly similar results (Figures S14, S15 & S13B). Average correlations between the hypothesis-driven and data-driven predictive-feature matrices were r = 0.99 (cognition), 0.84 (personality) and 0.92 (mental health).

### Control analyses

We performed several additional control analyses to ensure robustness of our results. First, we regressed age and sex (in addition to FD/DVARS) from the behavioral variables before prediction, which only decreased the prediction performance slightly (Figure S16).

Second, instead of multi-kernel FC prediction, we averaged functional connectivity across all brain states (Elliott *et al.* 2019) and utilized the resulting mean-FC for kernel regression. We found that mean-FC yielded worse prediction performance for cognition compared with multi-kernel regression (Figure S17, but not personality and mental health. This suggests that the improvement in predicting cognitive traits using multi-kernel FC was not simply due to more available data per individual.

Third, to ensure our results were robust to the regression model, we also performed linear ridge regression. We obtained similar prediction performance, but linear regression achieved worse COD (Figure S18). Remarkably, the feature-predictive matrices were highly similar for both linear regression and kernel regression (average r = 0.99), suggesting the predictive-feature matrices are robust to the choice of regression algorithm. We note that if we interpreted the regression weights directly without model inversion, then the agreement between kernel regression and linear regression “only” achieved an average correlation of r = 0.66. This observation confirms the importance of inverting the regression models (Haufe *et al.* 2014).

Fourth, we computed the predictive-feature matrices based on the single-kernel regression models and found that the results were highly similar to the predictive-feature matrices of the multi-kernel regression model (average r = 0.95).

## Discussion

In a large sample of typically developing children, we found that compared to resting-FC, task-FC of certain tasks improves prediction of cognition, but not (impulsivity-related) personality or mental health. Integrating resting-FC and task-FC further improves prediction of cognition and personality, but not mental health. By considering a large number of measures across cognition, personality and mental health, we found that these behavioral domains were predicted by distinct patterns of brain network features. However, within a behavioral domain (e.g., cognition) and across brain states, the predictive network features were similar, suggesting the potential existence of shared neural mechanisms explaining individual variation within each behavioral domain.

### Predictive brain network features cluster together within behavioral domains

Previous task-FC behavioral prediction studies have typically focused on specific cognitive traits, such as fluid intelligence (Greene *et al.* 2018), attention (Rosenberg *et al.* 2016) or reading comprehension (Gao *et al.* 2019). By exploring a wide range of behavioral measures, we gained insights into shared and unique predictive network features across traits within the same domain and across domains, as well as across brain states (rest and task). While there were differences among predictive network features within a behavioral domain (Figures S3-S6), the strong similarity was striking (Figures 5-6). This was especially the case for the cognitive domain (Figures 5-6 & S3-S6), where the average correlation of predictive network features across cognitive measures was 0.68.

Decades of studies, ranging from lesion to functional neuroimaging studies, have suggested the existence of brain networks that are specialized for specific cognitive functions (Petersen *et al.* 1988, Freiwald and Tsao 2010, Nomura *et al.* 2010, Laird *et al.* 2011, Yeo *et al.* 2015). For example, language tasks activate a specific network of brain regions (Binder *et al.* 1997, Fedorenko *et al.* 2012, Braga *et al.* 2019). Another example is the specific loss of episodic memory but not language after medial temporal lobe lesions (Scoville and Milner 1957, Corkin 2002). Of course, the networks that preferentially underpin aspects of behavior do not work in isolation and many studies have also emphasized information integration across specialized brain networks (van den Heuvel and Sporns 2011, Bzdok *et al.* 2016, Cohen and D’Esposito 2016, Bertolero *et al.* 2018). Lesion studies have also suggested that damage to connector hubs lead to deficits in multiple functional domains (Warren *et al.* 2014). Thus, while we did not expect predictive network features to be completely different across cognitive measures, we did not anticipate such strong similarity.

Similarly, in the case of mental health measures, while diagnostically distinct psychiatric disorders are likely the result of differentially disrupted brain systems, there is significant comorbidity among disorders and overlap in clinical symptoms (Kessler *et al.* 2011, Tamminga *et al.* 2013, Russo *et al.* 2014). Certain brain circuits have also been disproportionately reported to be transdiagnostically aberrant across multiple psychiatric and neurological disorders (Menon 2011, Whitfield-Gabrieli and Ford 2012, Goodkind *et al.* 2015, Baker *et al.* 2019, Kebets *et al.* 2019). For instance, there is evidence for the core role of frontoparietal network disruptions across psychiatric diagnosis (Cole, Repovš, *et al.* 2014). Therefore, similarly to cognition, we did not expect predictive network features to be completely different across mental health measures, but the degree of similarity was still surprising. These findings underscore the importance of studying multiple facets of psychopathology at once in order to better characterize covariation among symptoms to redefine psychiatric nosologies (Kozak and Cuthbert 2016, Kotov *et al.* 2017).

One possibility is that even though the regression models were trained on specific behavioral measures, the learned models might be predicting a broad behavior rather than the specific behaviors they were trained on. For example, in the case of cognition, perhaps the network features were simply predicting the *g* factor, a general cognitive ability that can account for half of the variance of cognitive test scores (Carroll 2003). In the case of mental health, the network features might be predicting the *p* factor, a general psychopathology factor that reflects individuals’ susceptibility to develop psychopathologies (Caspi *et al.* 2014). The similarity in predictive network features across the personality measures was less surprising since the personality measures we considered were mostly impulsivity-related. Thus, the regression models might simply be predicting an overall impulsivity trait (Leshem and Glicksohn 2007).

### Distinct brain network features support the prediction of cognition, personality, and mental health

We found that cognitive performance was predicted by a distributed set of network features across the whole brain with connectivity of salience and somatomotor networks being particularly notable (Figures 6C & 6D). The involvement of the salience network might not be surprising given its involvement in saliency, switching, attention and control (Menon and Uddin 2010). The prominent role of the somatomotor network was more surprising, although somatomotor regions have been reported to be associated with fluid intelligence (Greene *et al.* 2018), attention (Rosenberg *et al.* 2016), and general cognitive dysfunction (Kebets *et al.* 2019).

Similarly to cognitive performance, (impulsivity-related) personality measures were predicted by a distributed set of network features across the whole brain. In the case of personality, connectivity involving default and dorsal attentional networks was particularly prominent. While classical models of impulsivity have typically highlighted dysregulation in fronto-striatal circuits, these have been predominantly informed by animal lesion, PET and case-control task activation studies (Jentsch and Taylor 1999, Dalley *et al.* 2008, Beck *et al.* 2009, Buckholtz *et al.* 2010, Fineberg *et al.* 2010, Balodis *et al.* 2012, Cubillo *et al.* 2012). Conversely, fMRI studies of healthy participants have reported correlations between trait impulsivity and resting-FC in default (Inuggi *et al.* 2014, Golchert *et al.* 2017) and attentional (Golchert *et al.* 2017) networks. FC measured during the SST in attentional regions has also been found to be correlated with impulsivity in adults (Farr *et al.* 2012). Our whole-brain connectome approach not only supports the roles of default and attentional networks in impulsivity from these seed-based studies, but also extends these findings to children.

Finally, mental health measures were predicted by a distributed set of network features with the connectivity of default and frontoparietal control networks being particularly prominent. Connectivity involving default and frontoparietal regions have been linked to multiple psychiatric disorders and associated symptom profiles (Whitfield-Gabrieli and Ford 2012, Baker *et al.* 2014, 2019, Xia *et al.* 2018, Sha *et al.* 2019). We extend these findings by showing that the connectivity of these networks were important for predicting mental health in typically developing children prior to the onset of psychiatric illness and at a point where association cortices are still maturing.

### Resting and task network organization

A surprising result is that the predictive network features were similar across brain states (rest, MID, SST, N-Back) for all behavioral domains, particularly in the case of personality and mental health. On the one hand, task network reorganization has been shown to influence cognitive performance (Schultz and Cole 2016, Zuo *et al.* 2018). On the other hand, our results are consistent with studies showing that task states only modestly influence functional connectivity (Cole, Bassett, *et al.* 2014, Krienen *et al.* 2014, Bzdok *et al.* 2015) with inter-individual differences dominating task modulation (Gratton *et al.* 2018).

We note that a previous study (Gao *et al.* 2019) suggested that the regression models utilized different network features for prediction across different brain states, while another study (Greene *et al.* 2018) suggested that there was substantial overlap in predictive network features across resting-FC and task-FC. These discrepancies might arise because the previous studies only interpreted the most salient edges selected for prediction, which might yield unstable results. Here, we followed the elegant approach of Haufe and colleagues (2014) to invert the prediction models, leading to highly consistent predictive network features across two regression models (kernel regression and linear regression). A lack of inversion leads to weaker agreement between the two models.

Consistent with previous studies (Greene *et al.* 2018, Yoo *et al.* 2018, Fong *et al.* 2019), we found that task-FC outperforms resting-FC for the prediction of cognitive performance, at least in the case of N-back and MID. Although resting-FC was better than SST-FC for predicting cognition (Figure 2), we note that there was more resting-fMRI data than SST-fMRI data, which might explain the gap in performance. Here, we did not control for fMRI duration because our goal was to maximize prediction performance and to quantitatively characterize the predictive network features (Bzdok and Ioannidis 2019). Similarly, the prediction improvement from integrating information across brain states (multi-kernel regression) partly comes from the use of more fMRI data per child, but at least in the case of cognition, the improvement was not entirely due to more data (Figure S17).

Consistent with previous studies (Elliott *et al.* 2019, Gao *et al.* 2019, Jiang *et al.* 2019), we found that combining rest-FC and task-FC improved prediction of cognition. Extending upon this work, we demonstrate that combining rest-FC and task-FC modestly improved prediction of personality, but not mental health. We also found that regardless of using resting-FC, task-FC, or both resting-FC and task-FC, greater performance was achieved for predicting cognition than personality or mental health. This is again consistent with previous studies relating resting-fMRI with inter-individual variation in multiple behavioral domains (Dubois *et al.* 2018, Kong *et al.* 2019, Liégeois *et al.* 2019, Maglanoc *et al.* 2019).

### Strengths and limitations

One strength of our study was the use of a whole-brain connectomics approach to predict a wide range of behavioral traits. Many neurodevelopmental studies have focused on specific brain circuits (Bjork *et al.* 2004, Galvan *et al.* 2006, Van Leijenhorst *et al.* 2010, Satterthwaite *et al.* 2012, Gee *et al.* 2013, Swartz *et al.* 2014, Jalbrzikowski *et al.* 2017, Silvers *et al.* 2017). Yet, the human brain comprises functional modules that interact as a unified whole to support behavior (Spreng et al. 2010, Bertolero et al. 2015, Bassett and Sporns 2017). Therefore, whole-brain network-level approaches could provide critical insights into neurodevelopment that might be missed by studies focusing on specific networks. Our results were also robust across brain states, simple and more advanced predictive algorithms and recruitment sites. However, since the ABCD cohort comprises typically developing children, it is unclear how our results, especially those pertaining to mental health, might generalize to groups with clinical diagnoses. Furthermore, the cross-sectional nature of our study and the limited age range of the participants prevented us from thoroughly examining neurodevelopmental changes across time or age. Whole-brain neurodevelopmental studies have shown that functional networks become more distributed throughout adolescence (Fair *et al.* 2009, Supekar *et al.* 2009, Power *et al.* 2010). As such, it remains to be seen how the predictive network features from our study might be similarly affected by the developmental process. Lastly, we did not include any non-imaging features, which could have enriched our predictive models (Eickhoff and Langner 2019).

### Conclusions

Our study demonstrated that combining task-FC and resting-FC can yield improved predictions of cognition and personality, but not mental health. Each behavioral domain was predicted by unique patterns of brain network features that were distinct from other behavioral domains. These features were robust across brain states and regression approaches. Overall, our findings revealed distinct brain network features that account for individual variation across broad domains of behavior, yet are shared for behaviors within the same domain.

## Methods

### Participants

We considered data from 11875 children from the ABCD 2.0.1 release. After strict preprocessing quality control (QC) and considering only participants with complete rest-fMRI, task-fMRI and behavioral data, our main analyses utilized 1858 unrelated children (Figure 1A). See further details below.

### Imaging acquisition & processing

Images were acquired across 21 sites in the United States with harmonized imaging protocols for GE, Philips, and Siemens scanners. We used structural T1, resting-fMRI, and task-fMRI from three tasks: monetary incentive delay (MID), N-Back, stop signal task (SST). See Supplemental Methods S1 for details.

Minimally preprocessed T1 data were used (Hagler *et al.* 2019). The structural data were further processed using FreeSurfer 5.3.0 (Dale *et al.* 1999, Fischl, Sereno, and Dale 1999, Fischl, Sereno, Tootell, *et al.* 1999, Fischl *et al.* 2001, Ségonne *et al.* 2004, 2007), which generated accurate cortical surface meshes for each individual. Individuals’ cortical surface meshes were registered to a common spherical coordinate system (Fischl, Sereno, and Dale 1999, Fischl, Sereno, Tootell, *et al.* 1999). Individuals who did not pass recon-all QC (Hagler *et al.* 2019) were removed.

Minimally preprocessed fMRI data (Hagler *et al.* 2019) were further processed with the following steps: (1) removal of the first four frames, (2) slice time correction with the FSL library (Jenkinson *et al.* 2002, Smith *et al.* 2004), (3) motion correction using rigid body translation and rotation with FSL, and (4) alignment with the T1 images using boundary-based registration (Greve and Fischl 2009) with FsFast (http://surfer.nmr.mgh.harvard.edu/fswiki/FsFast). Functional runs with boundary-based registration costs greater than 0.6 were excluded. Framewise displacement (FD) (Jenkinson *et al.* 2002) and voxel-wise differentiated signal variance (DVARS) (Power *et al.* 2012) were computed using fsl_motion_outliers. Volumes with FD > 0.3 mm or DVARS > 50, along with one volume before and two volumes after, were marked as outliers and subsequently censored. Uncensored segments of data containing fewer than five contiguous volumes were also censored (Gordon *et al.* 2016, Kong *et al.* 2019). fMRI runs with over half of their volumes censored were removed. We also excluded individuals who did not have at least 4 minutes for each fMRI state (rest, MID, N-Back, SST) from further analysis.

The following nuisance covariates were regressed out of the fMRI time series: global signal, six motion correction parameters, averaged ventricular signal, averaged white matter signal, and their temporal derivatives (18 regressors in total). Regression coefficients were estimated from the non-censored volumes. We chose to regress the global signal because we were interested in behavioral prediction and global signal regression has been shown to improve behavioral prediction performance (Greene *et al.* 2018, Li *et al.* 2019). The brain scans were interpolated across censored frames using least squares spectral estimation (Power *et al.* 2014), band-pass filtered (0.009 Hz ≤ f ≤ 0.08 Hz), and projected onto FreeSurfer fsaverage6 surface space and smoothed using a 6 mm full-width half maximum kernel.

### Functional connectivity

We used a whole-brain parcellation comprising 400 cortical regions of interest (ROIs) (Schaefer *et al.* 2018) (Figure 1B) and 19 subcortical ROIs (Fischl *et al.* 2002) (Figure 1C). For each participant and each fMRI run, functional connectivity (FC) was computed as Pearson’s correlations between the average time series of each pair of ROIs. FC matrices were averaged across runs from each state, yielding a 419 × 419 FC matrix for each fMRI state (rest, MID, N-back, SST). Censored frames were ignored when computing FC.

### Behavioral data

We analyzed data from all available dimensional neurocognitive (Luciana *et al.* 2018) and mental health (Barch *et al.* 2018) assessments, yielding 16 cognitive, 9 mental health and 11 impulsivity-related personality measures. See Supplemental Methods S2 for more details. Participants who do not have all behavioral measures were excluded from further analysis.

### Single fMRI-state prediction

We used kernel ridge regression to predict each behavioral measure based on resting-FC, MID-FC, N-back-FC and SST-FC separately. We chose kernel regression because of its strong prediction performance in resting-FC based behavioral prediction (He *et al.* 2020). Briefly, let *y_i_* and *FC_i_* be the behavioral measure and FC of training individual*i*. Let *y_t_* and *FC_t_* be the behavioral measure and FC of a test individual. Then, kernel regression would predict the test individual’s behavior as the weighted average of the training individuals’ behavior, i.e., *ŷ_t_* ≈ ∑_*i*∈*training set*_ *Similarity*(*FC_i_*, *FC_t_*)*y_i_*, where *Similarity*(*FC_i_*, *FC_t_*) was defined as the Pearson’s correlation between *FC_i_* and *FC_t_*. Thus, kernel regression assumed that individuals with more similar FC exhibit more similar behavior. To reduce overfitting, an l_2_-regularization term was included (Kong *et al.* 2019, Li *et al.* 2019, He *et al.* 2020). Details of this approach can be found elsewhere (Kong *et al.* 2019, Li *et al.* 2019, He *et al.* 2020).

Kernel regression was performed within an inner-loop (nested) cross-validation procedure. More specifically, there were 22 ABCD sites. As recommended by the ABCD consortium, individuals from Philips scanners were excluded due to incorrect pre-processing. Our final sample for the main analysis comprised 1858 children. To reduce sample size variability across sites, we combined sites together to create 10 “site-clusters”, each containing at least 150 individuals (Table S4). Thus, participants within a site are in the same site-cluster.

We performed leave-3-site-clusters-out nested cross-validation for each behavioral measure with 120 replications. For each fold, a different set of 3 site-clusters was chosen as the test set. Kernel ridge regression parameters were estimated from the remaining 7 site-clusters using cross-validation. For model selection, the regularization parameter was estimated within the “inner-loop” of the inner-loop (nested) cross-validation procedure. For model evaluation, the trained kernel regression model was applied to all unseen participants from the test site-clusters.

Head motion (mean FD and DVARS) were regressed from each behavioral measure before the cross-validation procedure. More specifically, regression coefficients were estimated from the 7 training site-clusters and applied to the 3 test site-clusters. This regression procedure was repeated for each split of the data into 7 training site-clusters and 3 test site-clusters.

Prediction performance was measured by correlating predicted and actual measures (Finn *et al.* 2015). We also computed coefficients of determinations, which yielded similar conclusions.

### Multi-state prediction

To explore whether combining resting-FC and task-FC would result in better prediction accuracy, we utilized FC matrices from all four brain states (Rest, MID, SST, N-back) for prediction using a multi-kernel framework (Supplemental Methods S3). Similarly to single-kernel regression, multi-kernel regression assumed that subjects with similar FC exhibit similar behavioral scores. However, instead of taking into account FC from one fMRI state, here we utilized FC from all four fMRI states.

### Statistical tests of prediction accuracy

To test whether a model achieved better-than-chance accuracy, we performed permutation tests by shuffling behavioral measures across participants and repeating the entire leave-3-site-clusters-out nested cross-validation procedure. To compare two models, a permutation test was not valid, so the corrected resampled t-test was utilized (Nadeau and Bengio 2003, Bouckaert and Frank 2004). The resampled t-test corrected for the fact that accuracies of test folds were not independent. We corrected for multiple comparisons using FDR (q < 0.05).

### Model interpretation

As can be seen, multi-kernel FC yielded the best prediction performance. Models estimated for prediction can be challenging to interpret (Bzdok and Ioannidis 2019). Here, we utilized the approach from Haufe and colleagues (2014), yielding a 419 × 419 predictive-feature matrix for each FC state and each behavioral measure (Supplemental S4). A positive (or negative) predictive-feature value indicates that higher FC was associated with predicting greater (or lower) behavioral values.

The predictive-feature matrices were more similar among behavioral measures within the same behavioral domain (cognition, mental health and personality) than across domains. Thus, we averaged the predictive-feature matrices within the same behavioral domain (cognitive, mental health and personality) considering only behavioral measures that were successfully predicted by multi-kernel FC regression. This yielded a 419 × 419 predictive-feature matrix for each fMRI state and each behavioral domain.

Statistical significance of the predictive-feature values was tested using a permutation test (2000 permutations). To limit the number of multiple comparisons, tests were performed for each within-network and between-network block by averaging predictive-feature values within and between 18 networks (Figures 6B & 6C). We corrected for multiple comparisons using FDR (q < 0.05).

### Control analyses

Because the multi-kernel model contained more input data compared to the single-kernel models, we explored the potential effect of the amount of input data on model performance. To this end, we performed a single-kernel ridge regression on a general functional connectivity matrix created by averaging the functional connectivity across all fMRI conditions (rest + MID + N-Back + SST) to predict behaviors, which we called Mean FC. We then compared the performance of the Mean FC model with the best single-kernel fMRI model (e.g. N-Back only) and the multi-kernel model. To assess the impact of age and sex on model performance, we performed kernel ridge regression to predict behaviors after regressing out age and sex, in addition to head motion (mean FD and DVARS).

## Data availability

The ABCD data are publicly available: http://dx.doi.org/10.15154/1504041.

## Code availability

Preprocessing utilized code from previously published pipelines (Kong *et al.* 2019, Li *et al.* 2019):

https://github.com/ThomasYeoLab/CBIG/tree/master/stable_projects/preprocessing/CBIG_fMRI_Preproc2016 Preprocessing code specific to this study can be found here: GITHUB_LINK. Analysis code specific to this study can be found here: GITHUB_LINK.

## Supplementary methods and materials

### S1. MRI acquisition

For each participant, twenty minutes of resting-state fMRI data were acquired in four 5-minute runs. The task fMRI data consisted of three tasks (MID, N-back, SST) that were each acquired over two runs (for a total of six task fMRI runs). Each fMRI run was acquired in 2.4 mm isotropic resolution with a TR of 800 ms. The structural data consisted of one 1 mm isotropic scan for each participant. Full details of image acquisition can be found elsewhere (Casey *et al.* 2018).

### S2. Behavioral data

We analyzed data from all available dimensional neurocognitive (Luciana *et al.* 2018) and mental health (Barch *et al.* 2018) assessments. For the neurocognitive assessments, we included the NIH Toolbox, Rey Auditory Verbal Learning Test, Little Man Task, and the matrix reasoning subscale from the Wechsler Intelligence Scale for Children-V, in order to measure different aspects of cognition. For the mental health assessments, we included the Achenbach Child Behavior Check List (CBCL), the mania scale from the Parent General Behavior Inventory, Pediatric Psychosis Questionnaire. For the personality measures, we included the Modified UPPS-P for Children and Behavioral Inhibition and Activation scales. See Tables S1 & S2 for more details for each individual scale.

**Table S1.** Behavioral measures used in this study.

| Scale | Subscale/Measure |
| --- | --- |
| NIH Toolbox (Hodes *et al.* 2013) | Flanker<br>List sorting working memory<br>Dimensional change card sort<br>Oral reading recognition<br>Pattern comparison processing speed<br>Picture sequence memory test<br>Picture vocabulary test<br>Cognition fluid composite<br>Crystallized composite<br>Cognition total composite |
| Rey Auditory Verbal Learning Test (RAVLT) (Strauss *et al.* 2006) | Short delay recall<br>Long delay recall |
| Little Man Task (Acker and Acker 1982) | Accuracy<br>Reaction time (correct responses)<br>Efficiency |
| Wechsler Intelligence Scale for Children-V (WISC-V) (Wechsler 2014) | Matrix reasoning |
| Achenbach Child Behavior Check List<br>(Achenbach and Rescorla 2013) | Anxious/Depressed<br>Withdrawn/Depressed<br>Somatic complaints<br>Social problems<br>Thought problems<br>Attention problems<br>Rule-breaking behavior<br>Aggressive behavior |
| Parent General Behavior Inventory<br>(Youngstrom *et al.* 2013) | Mania |
| Pediatric Psychosis Questionnaire - Brief<br>Version (Loewy *et al.* 2012) | Total number of psychosis symptoms<br>Symptom severity score |
| Modified UPPS-P for Children from PhenX<br>(Lynam 2013) | Negative urgency<br>Positive urgency<br>Lack of planning<br>Lack of perseverance<br>Sensation seeking |
| Behavioral Inhibition & Activation<br>(Pagliaccio *et al.* 2016) | Behavioral inhibition sum<br>Reward responsiveness<br>Drive<br>Fun seeking |

**Table S2.**
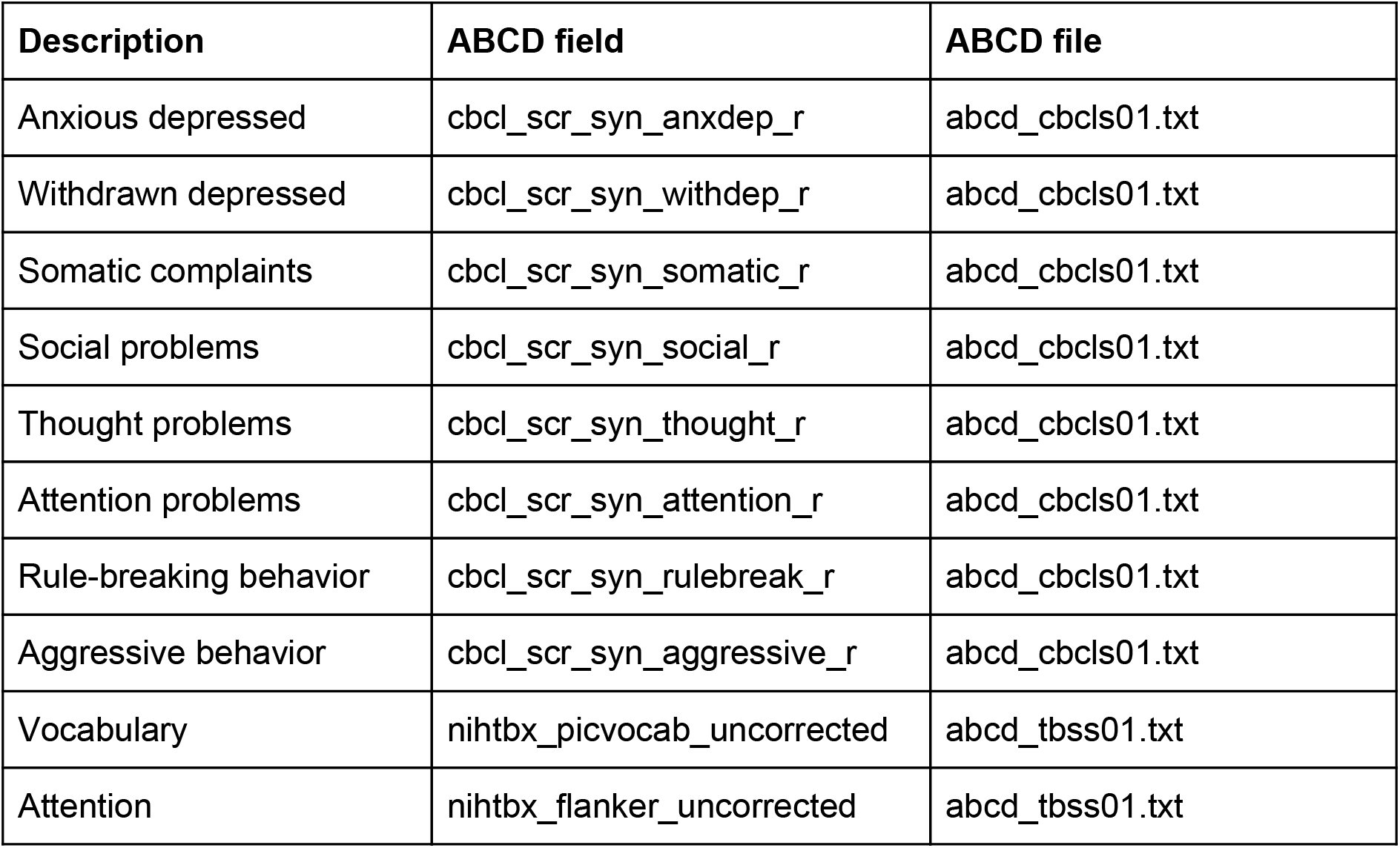
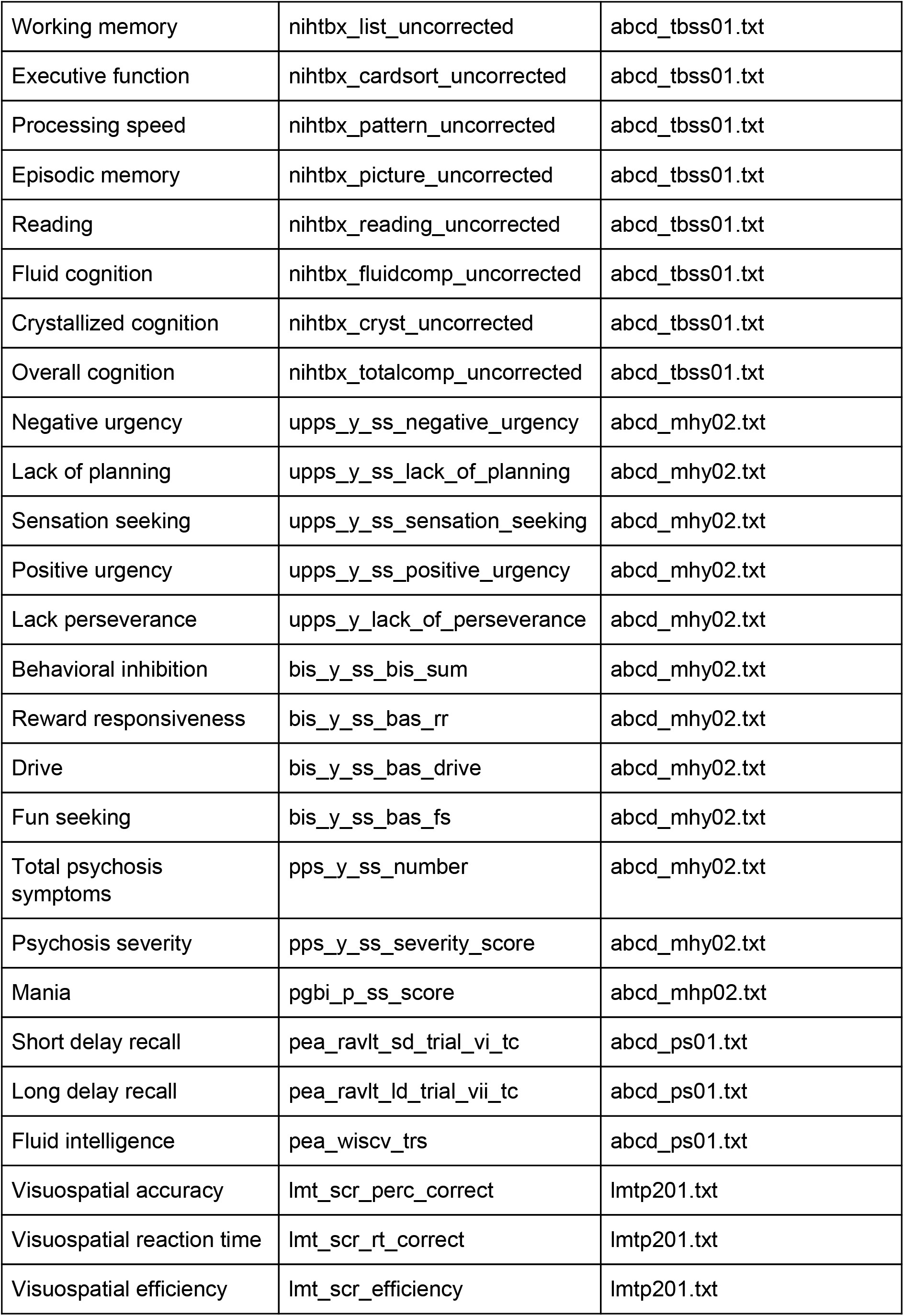
Lookup table showing the original ABCD variable names with the corresponding descriptive labels used in the manuscript. More details of the behavioral measures can be found in the ABCD data dictionary.

### S3. Multi-kernel ridge regression

#### S3.1. Single-kernel ridge regression

For completeness, we provide a brief explanation of single-kernel ridge regression. The following section is adapted from our previous study (Kong et al., 2019). Suppose we have *M* training subjects. Let *y_i_* be the behavioral measure (e.g., fluid intelligence) and *FC_i_* be the vectorized FC (considering only lower triangular matrix) of the *i*-th training subject. Given and, {*y*_1_, *y*_2_,⋯,*y_N_*} the {*FC*_1_, *FC*_2_,⋯,*FC_M_*} kernel regression model is written as:

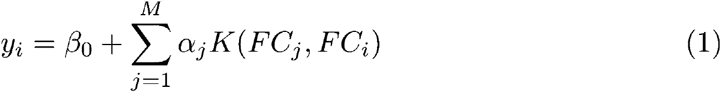

where *β*_0_ is the bias term and *K*(*FC_j_*, *FC_i_*) is the functional connectivity similarity between the *i*-th and *j*-th training subjects. *K*(*FC_j_*, *FC_i_*) is defined by the correlation between the vectorized FC of the two subjects. The choice of correlation is motivated by previous fingerprinting and behavioral prediction studies (Finn *et al.* 2015, Li *et al.* 2019, He *et al.* 2020).

To estimate ***α*** and *β*_0_ from the training set, let ***y*** = [*y*_1_, *y*_2_,⋯*y_M_*]^T^, ***α*** = [*α*_1_, *α*_2_,⋯*α_M_*]^T^ and 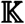 be the *M* × *M* kernel similarity matrix, whose (*j*, *i*)-th element is *K*(*FC_j_*, *FC_i_*). Note that we can rewrite Eq. (1) as 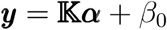. We can then estimate ***α*** and *β*_0_ by minimizing the following l_2_-regularized cost function:

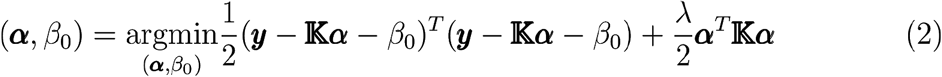

where *λ* controls the importance of the l_2_-regularization and is estimated within the inner-loop cross-validation procedure. We emphasize that the test set was not used to estimate *λ*. Once ***α*** and *β*_0_ have been estimated from the training set, the predicted behavior of test subject *t* is given by

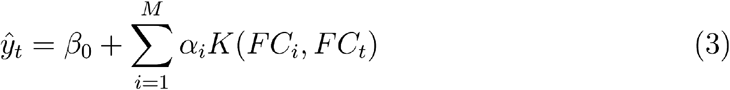

#### S3.2. Multi-kernel ridge regression

Single-kernel ridge regression uses data from a single fMRI brain state for prediction. To extend to multiple fMRI brain states, we can construct one kernel similarity matrix for each fMRI brain state. Suppose we have *M* training subjects and *R* fMRI brain states. Let *y_i_* be the behavioral measure of the *i*-th training subject. Let *FC_ir_* be the vectorized FC of the *i*-th training subject for the *r*-th fMRI brain state. The multi-kernel regression model can be written as:[

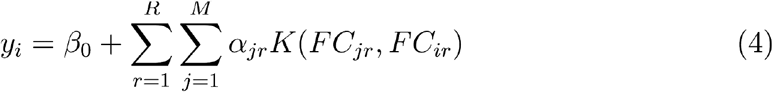

where *β*_0_ is the bias term and *K*(*FC_jr_*, *FC_ir_*) is the functional connectivity similarity between the *i*-th and *j*-th training subjects for the *r*-th brain state. Like before, is defined *K*(*FC_jr_*, *FC_ir_*) by the correlation between the vectorized FC of the two subjects for the *r*-th brain state.

Let ***y*** = [*y*_1_, *y*_2_,⋯*y_M_*]^T^ and ***α_r_*** = [*α*_1,*r*_, *α*_2,*r*_,⋯*α_M,*r*_*]^T^. Suppose 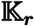 is the *M* × *M* kernel similarity matrix for the *r*-th brain state, whose (*j*, *i*)-th element is *K*(*FC_jr_*, *FC_ir_*). We can estimate ***α_r_*** and *β*_0_ by minimizing the following l_2_-regularized cost function:

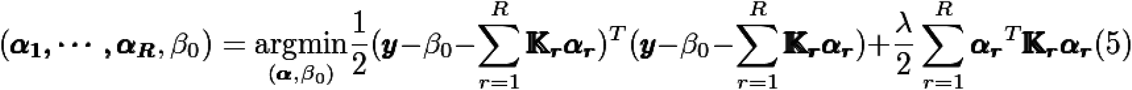

where *λ_r_* controls the importance of the l_2_-regularization for the *r*-th kernel. Here, *λ_r_* is estimated within the inner-loop cross-validation procedure using Gaussian-process optimization (Kawaguchi et al., 2015). We emphasize that the test set was not used to estimate *λ_r_*. Once ***α_r_*** and *β*_0_ have been estimated from the training set, the predicted behavior of test subject *t* is given by

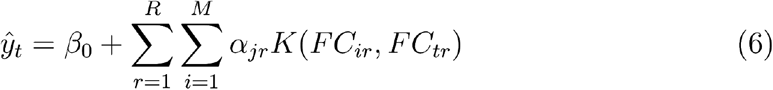

#### S3.3. Coefficient of determination (COD)

Suppose *N* is the number of test subjects, *y_t_* and *ŷ_t_* are the groundtruth and predicted behavior measure of the *t*-th test subject respectively, and 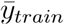 is the mean of the behavioral measure of all training subjects. The coefficient of determination is defined as follows:

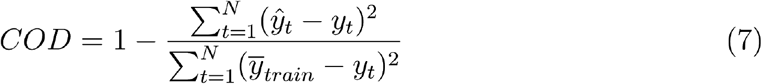

Thus, a larger COD indicates more accurate prediction. A negative value implies that we are better off using the mean behavior of the training subjects to predict the behavior of the test subject instead of using the FC data.

### S4. Predictive-feature matrices

To interpret which brain edges were important for the multi-kernel FC model, we utilized an elegant approach (Haufe *et al.* 2014) to invert the prediction model. Failure to invert the model leads to uninterpretable results (Haufe *et al.* 2014). Let us consider the functional connectivity between brain regions *a* and *b*. We would like to compute the predictive-feature value of the functional connection *p_ab_* for the multi-kernel FC model. A positive value (or negative) predictive-feature value for an edge, indicating that higher FC between brain regions *a* and *b* was associated with predicting greater (or lower) behavioral values.

Let *FC_ab_* be the normalized functional connectivity strength between brain regions *a* and *b* for all training subjects. Therefore, *FC_ab_* is an *N* × 1 vector where *N* is the number of training subjects. Normalization was performed so that the FC of each subject has zero mean and unit norm. Let *ŷ* be the prediction of the training subjects’ behavioral measure based on the estimated kernel regression model. Therefore *ŷ* is an *N* × 1 vector where *N* is the number of training subjects. According to Haufe and colleagues (2014), *p_ab_* = *covariance*(*FC_ab_*, *ŷ*).

However, because we would like to compare across different behavioral measures, the scale of *ŷ* is very different across behavioral measures. Thus, we computed *p_ab_* = *covariance*(*FC_ab_*, *ŷ*)/*variance*(*ŷ*), which does not change the relative predictive-feature values among edges, but allows for comparisons between behavioral measures. We note that the above formula is applied to the training set, because we want to interpret the trained model. However, recall that we performed leave-3-site-clusters-out nested cross-validation for each behavioral measure with 120 replications. Thus we computed the predictive-feature values for each replication and averaged across the 120 replications.

## Supplementary results

**Table S3.** Demographic information for included and excluded participants in ABCD 2.0.1

|  | Included | Excluded |
| --- | --- | --- |
| **N** | 1858 | 10017 |
| **Age in months (mean (s.d.))** | 120.34 (7.43) | 118.68 (7.44) |
| **Female (%)** | 1025 (55.17) | 4656 (46.48) |
| **Race/ethnicity (%)** |  |  |
| **Asian** | 55 (2.96) | 197 (1.97) |
| **Black** | 143 (7.70) | 1636 (16.33) |
| **Hispanic** | 324 (17.44) | 2083 (20.79) |
| **White** | 1145 (61.63) | 5029 (50.20) |
| **Other** | 187 (10.06) | 1058 (10.56) |
| **Unknown** | 4 (0.21) | 14 (0.14) |
| **Household income (%)** |  |  |
| **< 50 000** | 360 (19.38) | 2862 (28.57) |
| **≥ 50 000 & < 100 000** | 517 (27.82) | 2553 (25.49) |
| **≥ 100 000** | 873 (46.99) | 3692 (36.86) |
| **Unknown** | 108 (5.81) | 910 (9.08) |

**Table S4.** Distribution of the included sample (n=1858) by site and scanner

| **ABCD Site** | **Make** | **Model** | **N** | **Site-cluster** |
| --- | --- | --- | --- | --- |
| 16 | Siemens | Prisma | 292 | A |
| 13 | GE | Discovery MR750 | 161 | B |
| 4 | GE | Discovery MR750 | 145 | C |
| 22 | GE | Discovery MR750 | 12 | C |
| 14 | Siemens | Prisma/Prisma fit | 135 | D |
| 15 | Siemens | Prisma fit | 27 | D |
| 6 | Siemens | Prisma fit | 131 | E |
| 9 | Siemens | Prisma fit | 52 | E |
| 10 | GE | Discovery MR750 | 127 | F |
| 11 | Siemens | Prisma | 52 | F |
| 3 | Siemens | Prisma | 120 | G |
| 5 | Siemens | Prisma fit | 56 | G |
| 2 | Siemens | Prisma fit | 110 | H |
| 7 | Siemens | Prisma fit | 55 | H |
| 8 | GE | Discovery MR750 | 63 | I |
| 20 | Siemens | Prisma/Prisma fit | 92 | I |
| 12 | Siemens | Prisma fit | 79 | J |
| 18 | GE | Discovery MR750 | 73 | J |
| 21 | Siemens | Prisma fit/Prisma | 76 | J |

**Figure S1.**
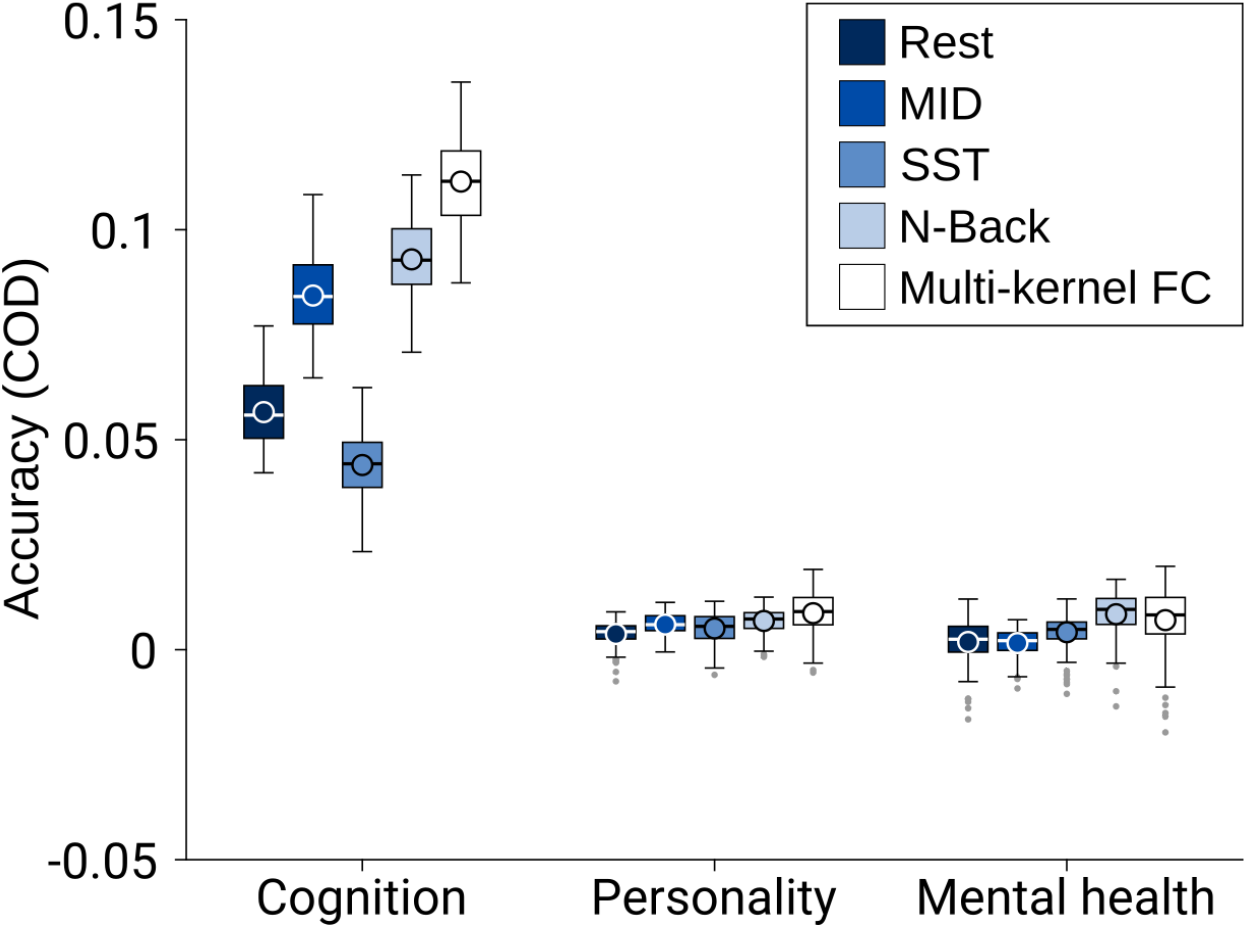
Cross-validated prediction performance (coefficient of determination; COD) using kernel ridge regression for resting-state and task-states (MID, SST, N-Back). Multi-kernel FC utilized FC from all 4 brain states for prediction. Higher COD indicates greater variance predicted relative to the mean of the training data.

**Figure S2.**
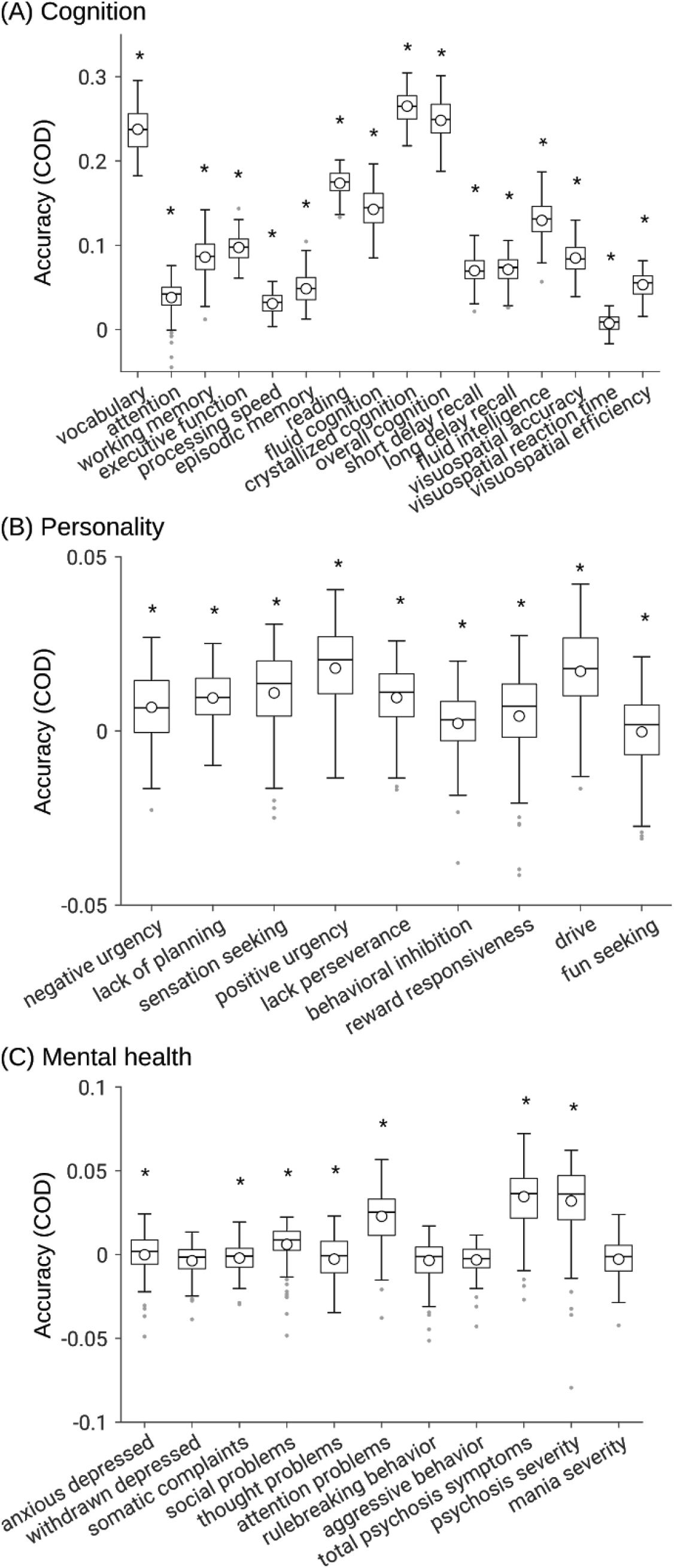
Cross-validated prediction performance (coefficient of determination; COD) using multi-kernel ridge regression by exploiting resting-FC, MID-FC, SST-FC and N-back-FC jointly. (A) Cognitive measures. (B) Personality measures. (C) Mental health measures. * denotes above chance prediction after correcting for multiple comparisons (FDR q < 0.05).

**Figure S3.**
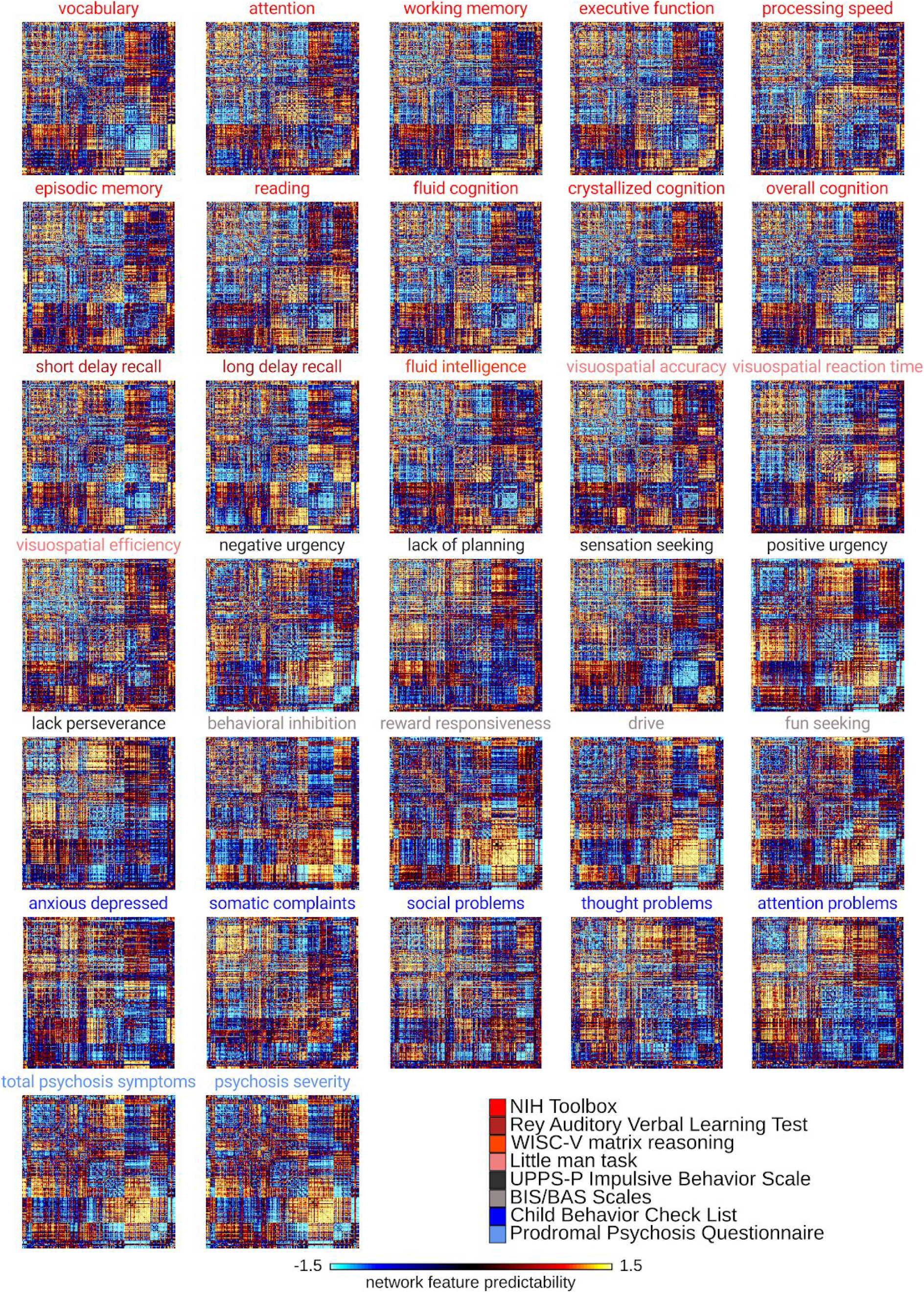
Resting-FC predictive-feature matrices for each significantly predicted behavior. For visualization, the values within each matrix were divided by their standard deviations.

**Figure S4.**
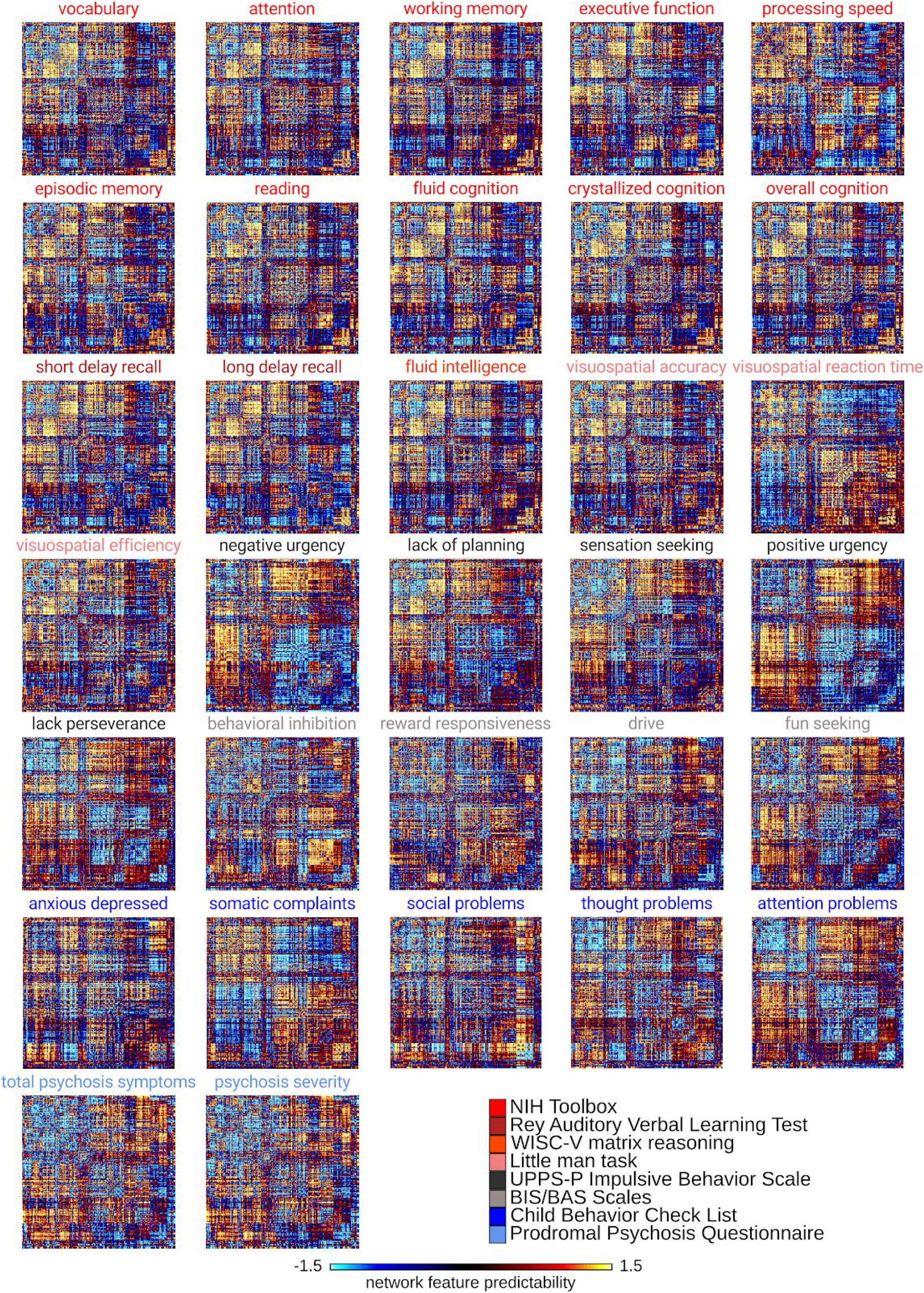
MID-FC predictive-feature matrices for each significantly predicted behavior. For visualization, the values within each matrix were divided by their standard deviations.

**Figure S5.**
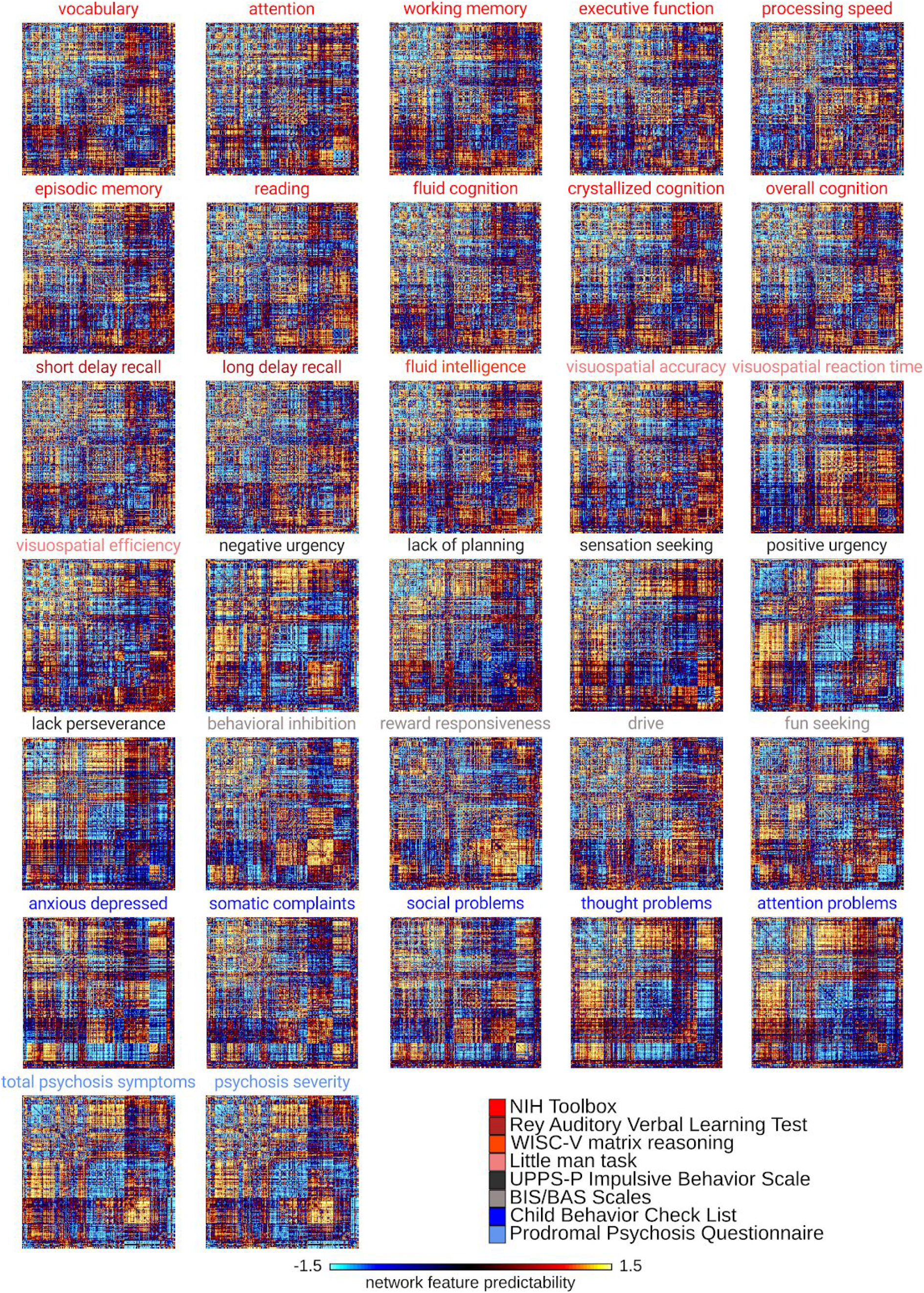
SST-FC predictive-feature matrices for each significantly predicted behavior. For visualization, the values within each matrix were divided by their standard deviations.

**Figure S6.**
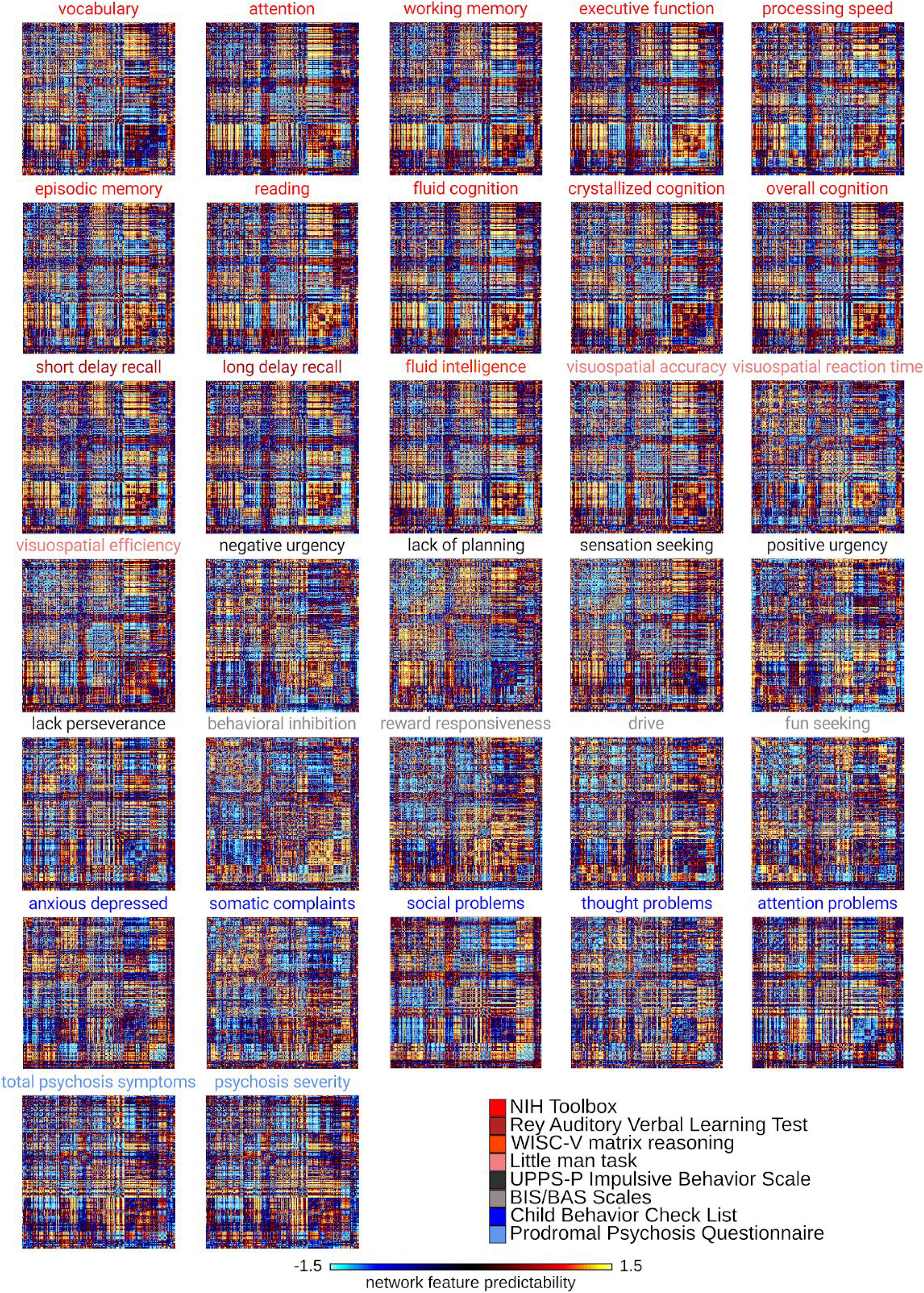
N-Back-FC predictive-feature matrices for each significantly predicted behavior. For visualization, the values within each matrix were divided by their standard deviations.

**Figure S7.**
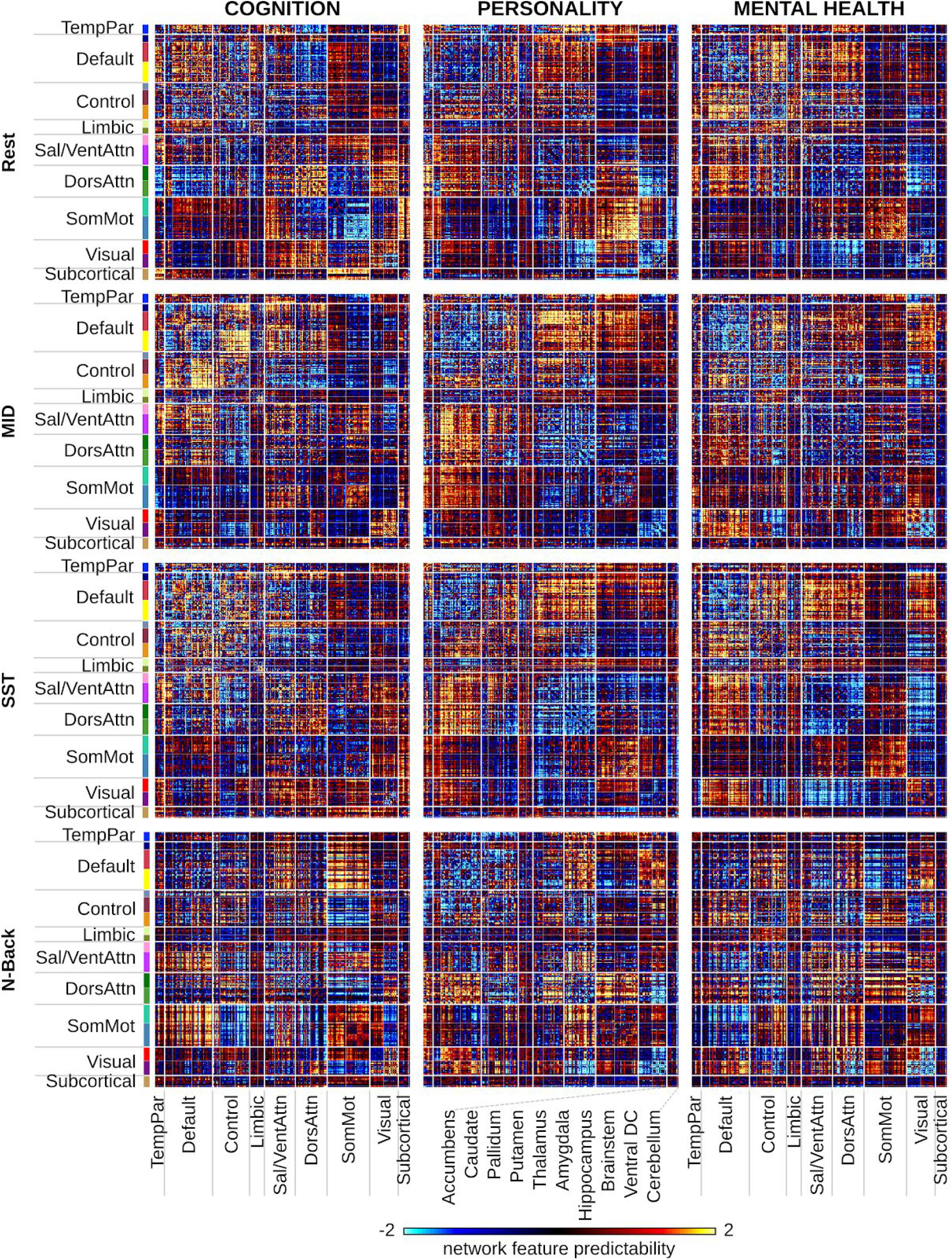
Predictive-feature matrices for each brain state (Rest, MID, SST, N-Back) averaged across all behavioral measures within each hypothesis-driven behavioral domain (cognition, personality, mental health). For visualization, the values within each matrix were divided by their standard deviations.

**Figure S8.**
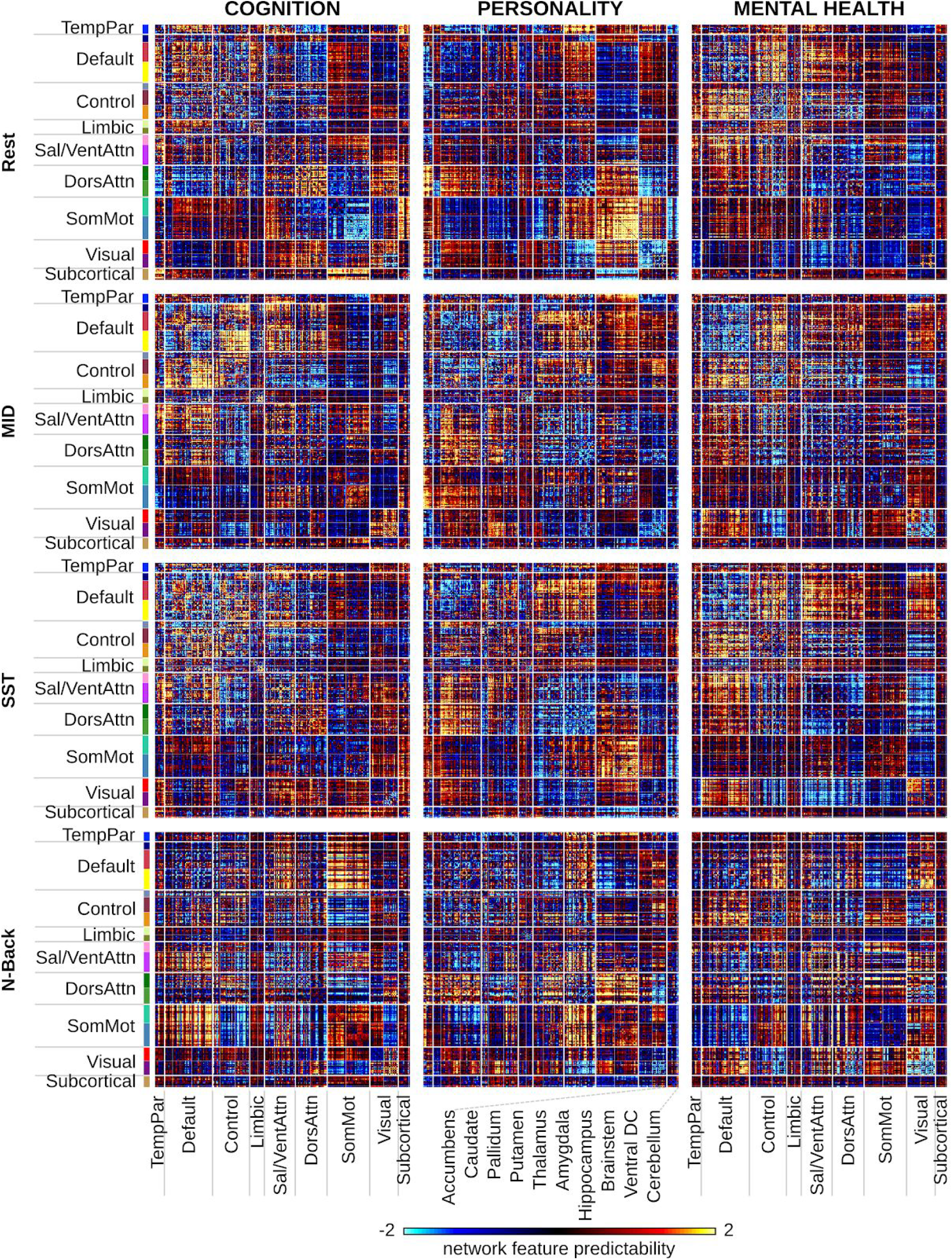
Predictive-feature matrices for each brain state (Rest, MID, SST, N-Back) averaged across all behavioral measures within each data-driven behavioral cluster (cognition, personality, mental health). For visualization, the values within each matrix were divided by their standard deviations.

**Figure S9.**
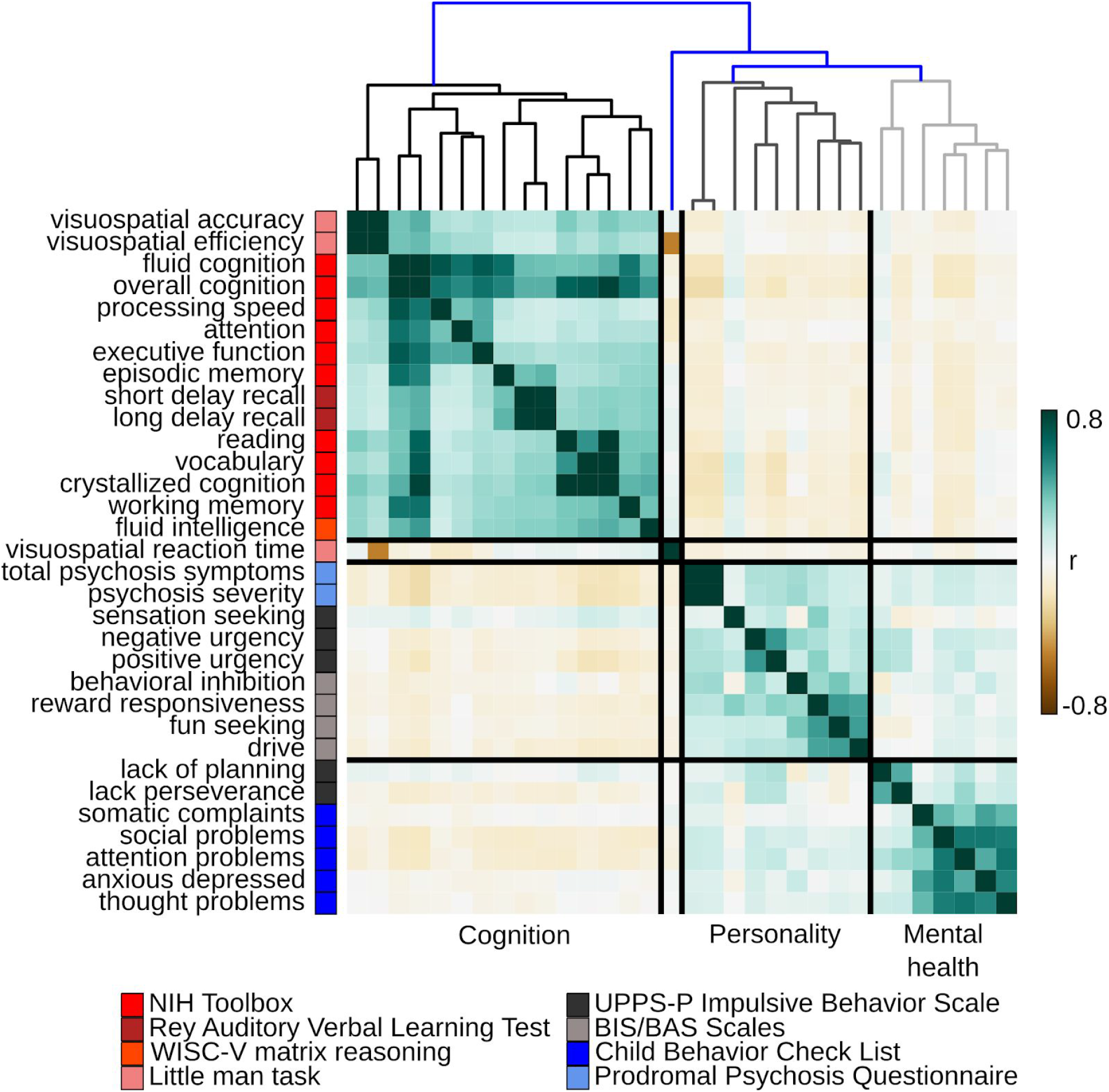
Hierarchical clustering of actual behavioral scores. Clustering was performed using hierarchical agglomerative average linkage (UPGMA) clustering as implemented in scipy 1.2.1 (Virtanen *et al.* 2020).

**Figure S10.**
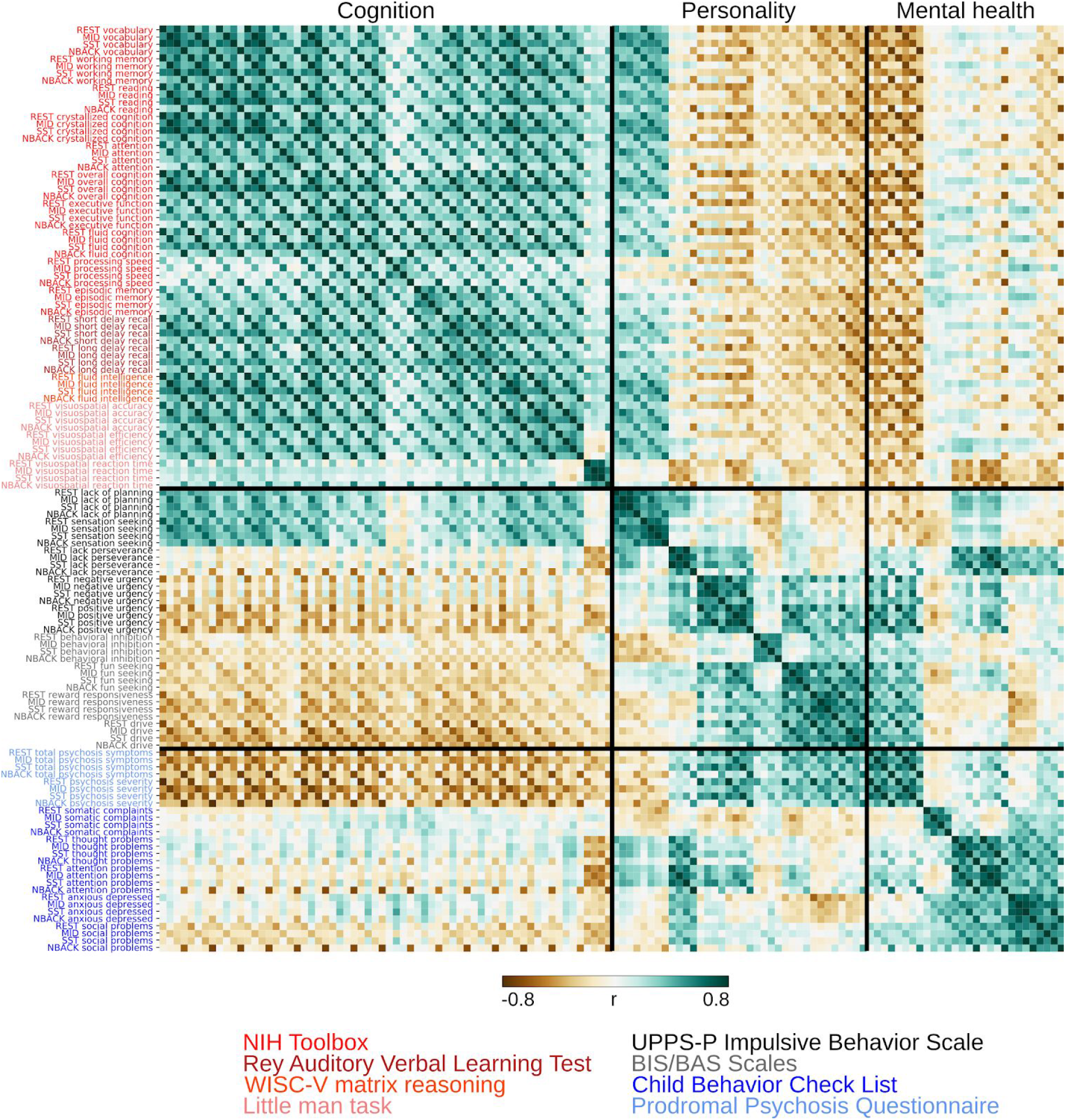
Similarity of predictive-network features for each significantly predicted behavior and brain state. The behavioral measures were ordered based on hypothesis-driven behavioral domains (cognition, personality and mental health). For each behavior, the brain states were ordered by Rest, MID, SST and finally N-Back. Red font indicates cognitive measures. Black/grey font indicates personality measures. Blue font indicates mental health measures. Predictive-network features were highly correlated within each hypothesis-driven behavioral domain and across brain states.

**Figure S11.**
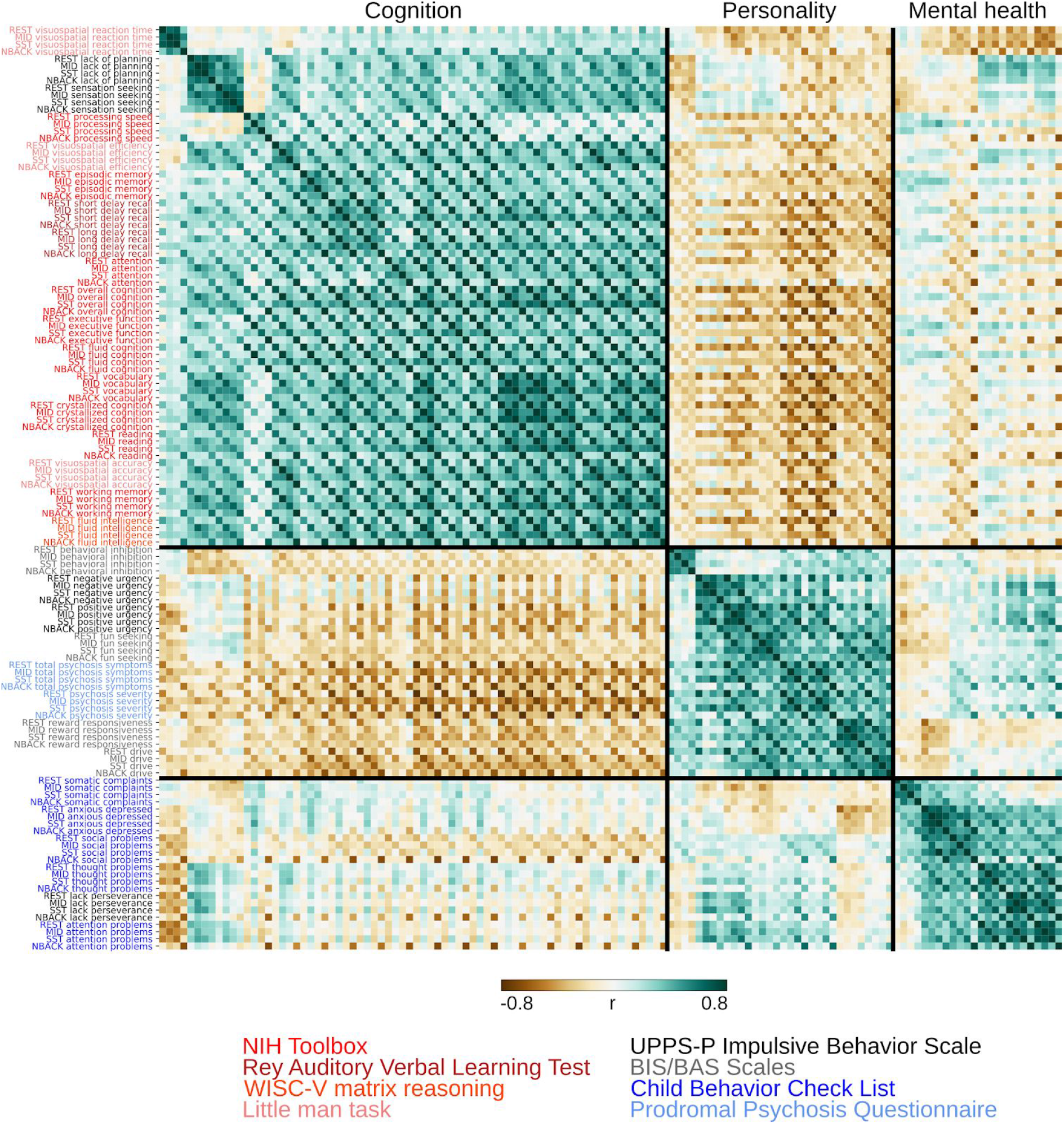
Similarity of predictive-network features for each significantly predicted behavior and brain state. The behavioral measures were ordered based on data-driven behavioral clusters (cognition, personality and mental health). For each behavior, the brain states were ordered by Rest, MID, SST and finally N-Back. Red font indicates cognitive measures. Black/grey font indicates personality measures. Blue font indicates mental health measures. Predictive-network features were highly correlated within each hypothesis-driven behavioral domain and across brain states.

**Figure S12.**
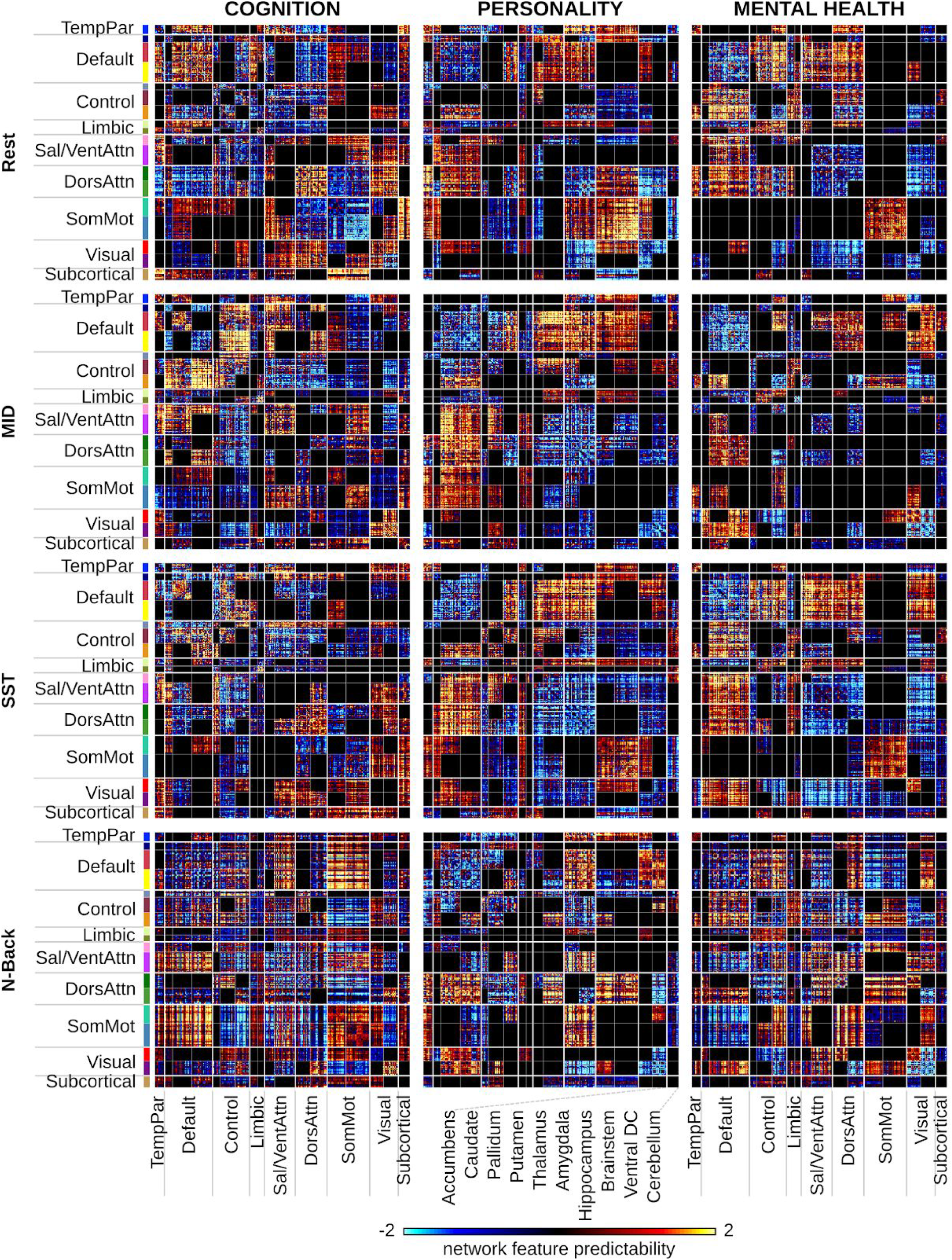
Predictive-feature matrices showing significant network blocks for each hypothesis-driven behavioral domain (cognitive, personality, mental health) for each brain state (Rest, MID, SST, N-Back) after permutation testing. For visualization, the values within each matrix were divided by their standard deviations.

**Figure S13.**
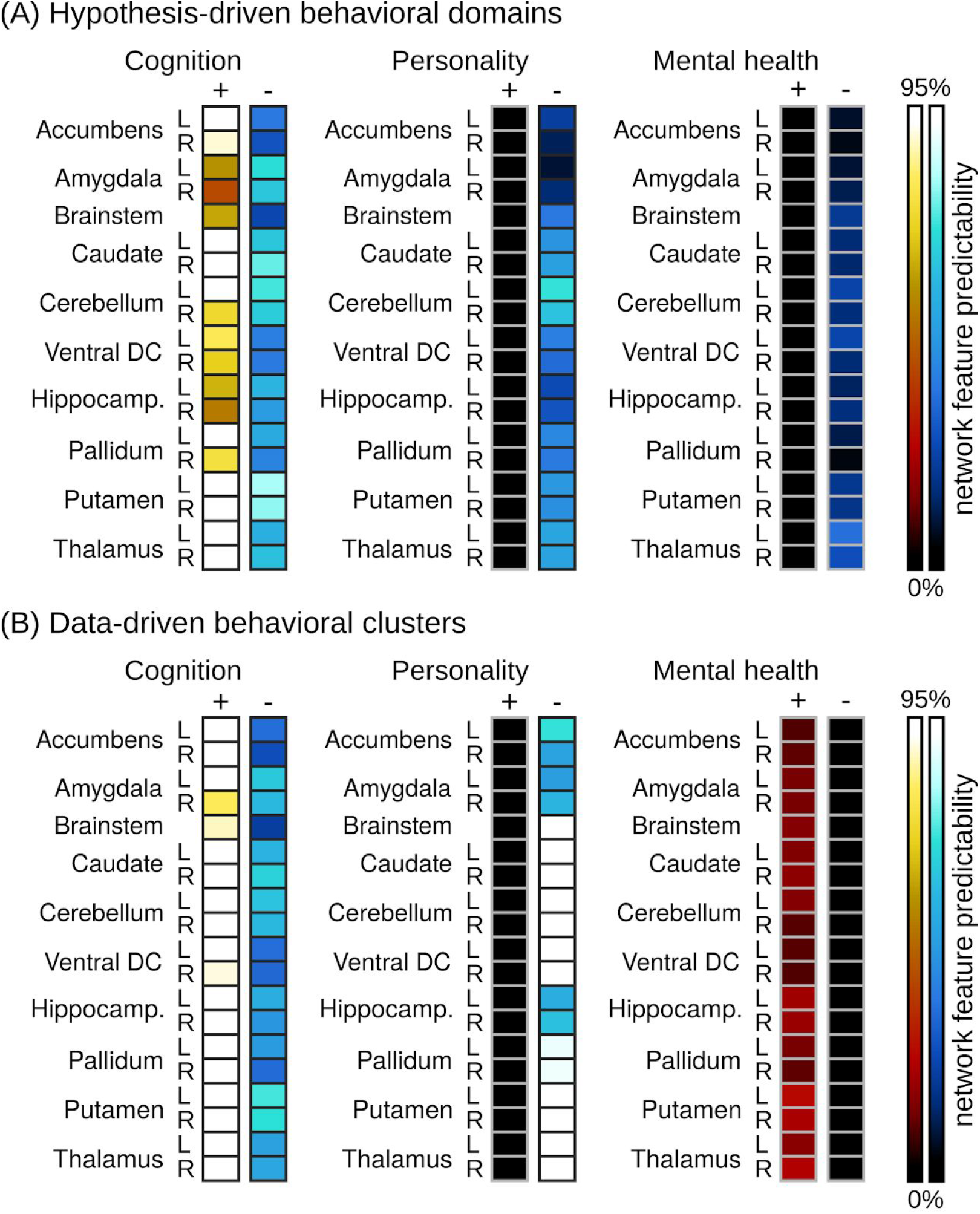
Heatmaps showing network feature predictability of each subcortical region for (A) each hypothesis-driven behavioral domain and (B) each data-driven behavioral cluster. See Figures 6C and 6D for the cortical maps of the hypothesis-driven behavioral domains and Figures S15C and S15D for the cortical maps of the data-driven behavioral clusters.

**Figure S14.**
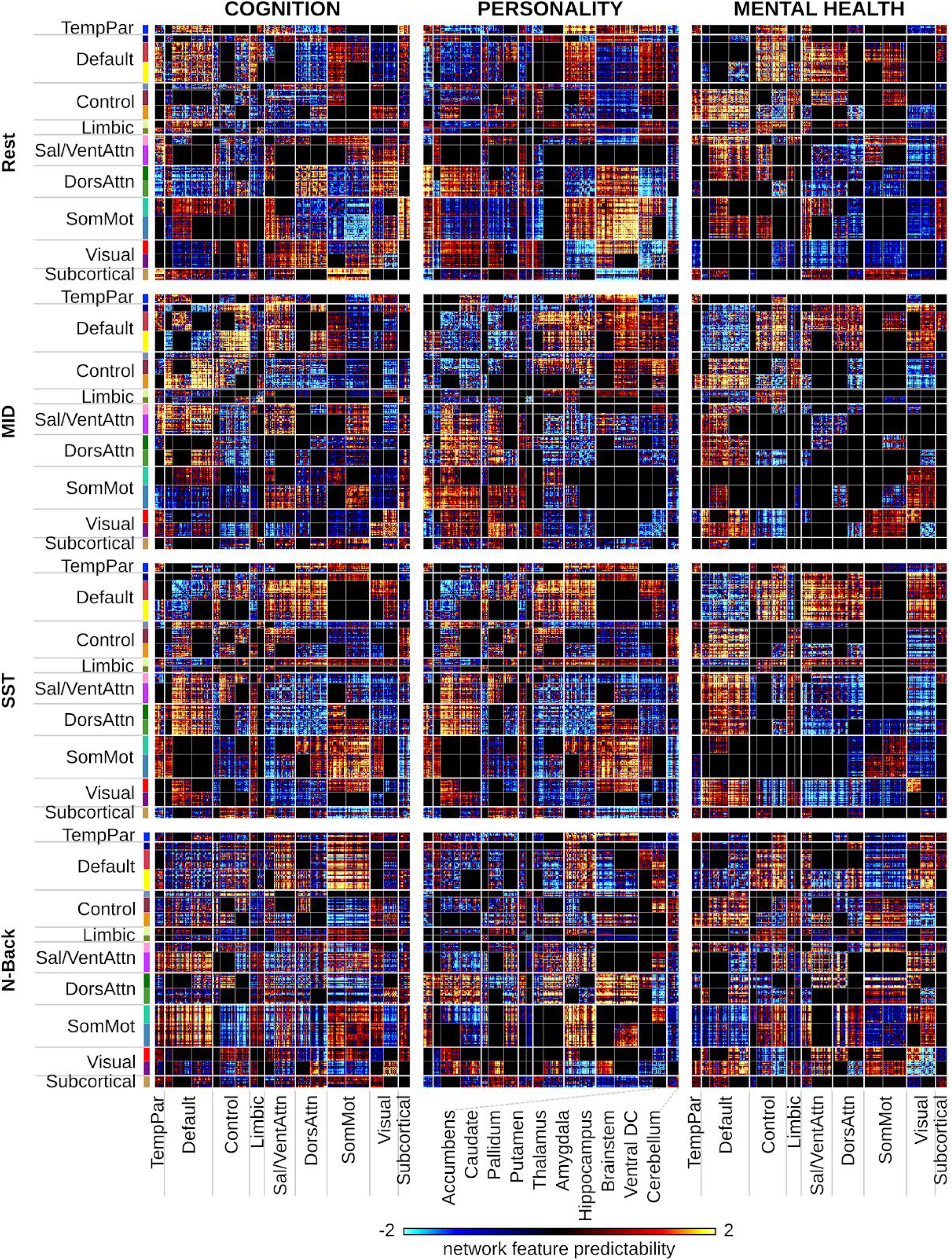
Predictive-feature matrices showing significant network blocks for each data-driven behavioral cluster (cognitive, personality, mental health) and for each brain state (Rest, MID, SST, N-Back) after permutation testing. For visualization, the values within each matrix were divided by their standard deviations.

**Figure S15.**
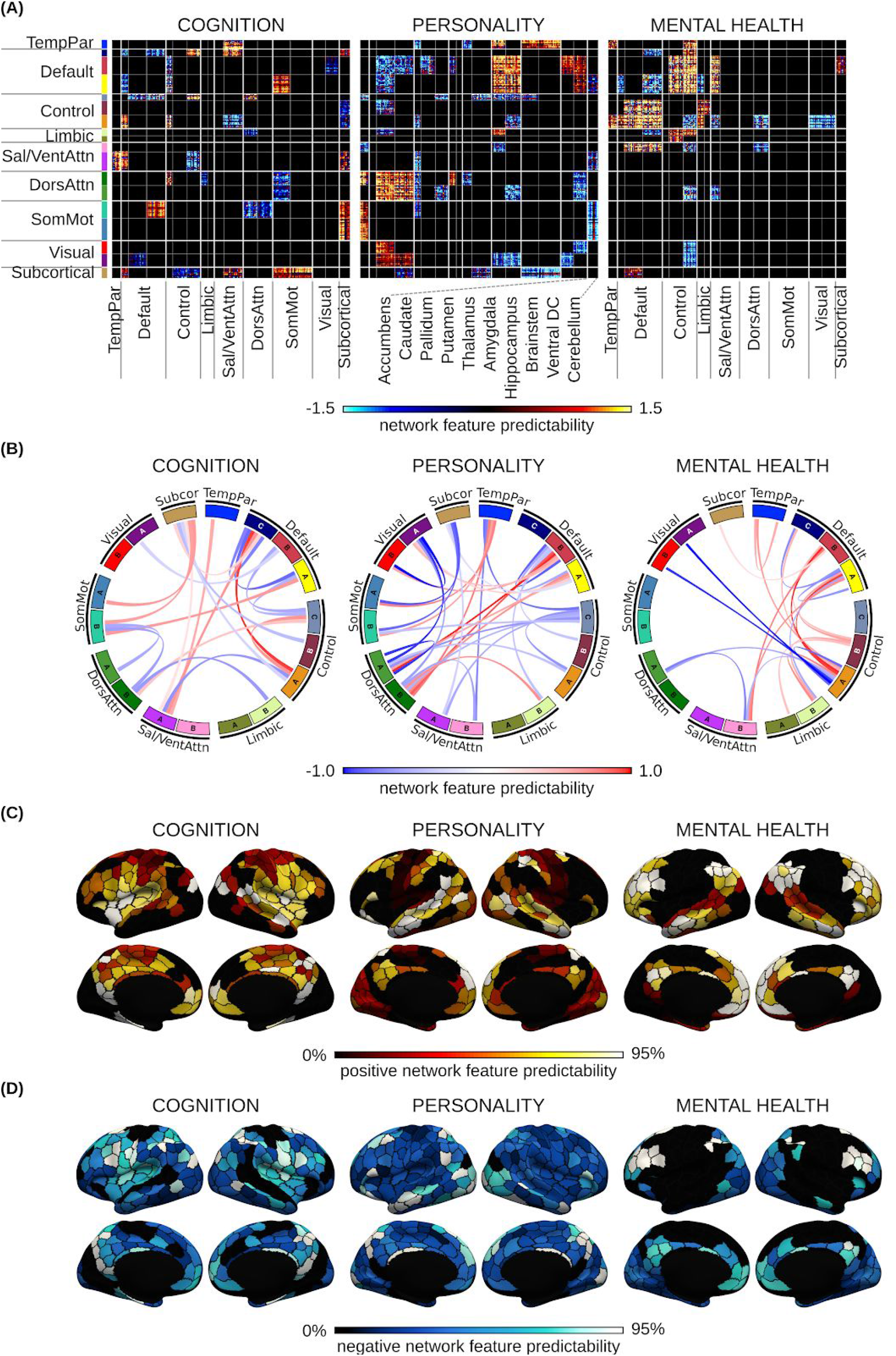
Predictive brain network features for predicting cognition, personality and mental health. This figure is the same as Figure 6 but using data-driven behavioral clusters, instead of hypothesis-driven behavioral domains. (A) Predictive-feature matrices averaged across brain states, considering only within-network and between-network blocks that were significant across all four brain states (Rest, MID, SST, N-Back). (B) Predictive network connections obtained by averaging the matrices in panel (A) within each between-network and within-network block. (C) Positive predictive features obtained by summing positive predictive-feature values across the rows of panel (A). A higher value for a brain region indicates that stronger connectivity yielded a higher prediction for the behavioral measure. (D) Negative predictive features obtained by summing negative predictive-feature values across the rows of panel (A). A higher value for a brain region indicates that weaker connectivity yielded a greater prediction for the behavioral measure. Conclusions were highly similar using hypothesis-driven behavioral domains (Figure 7).

**Figure S16.**
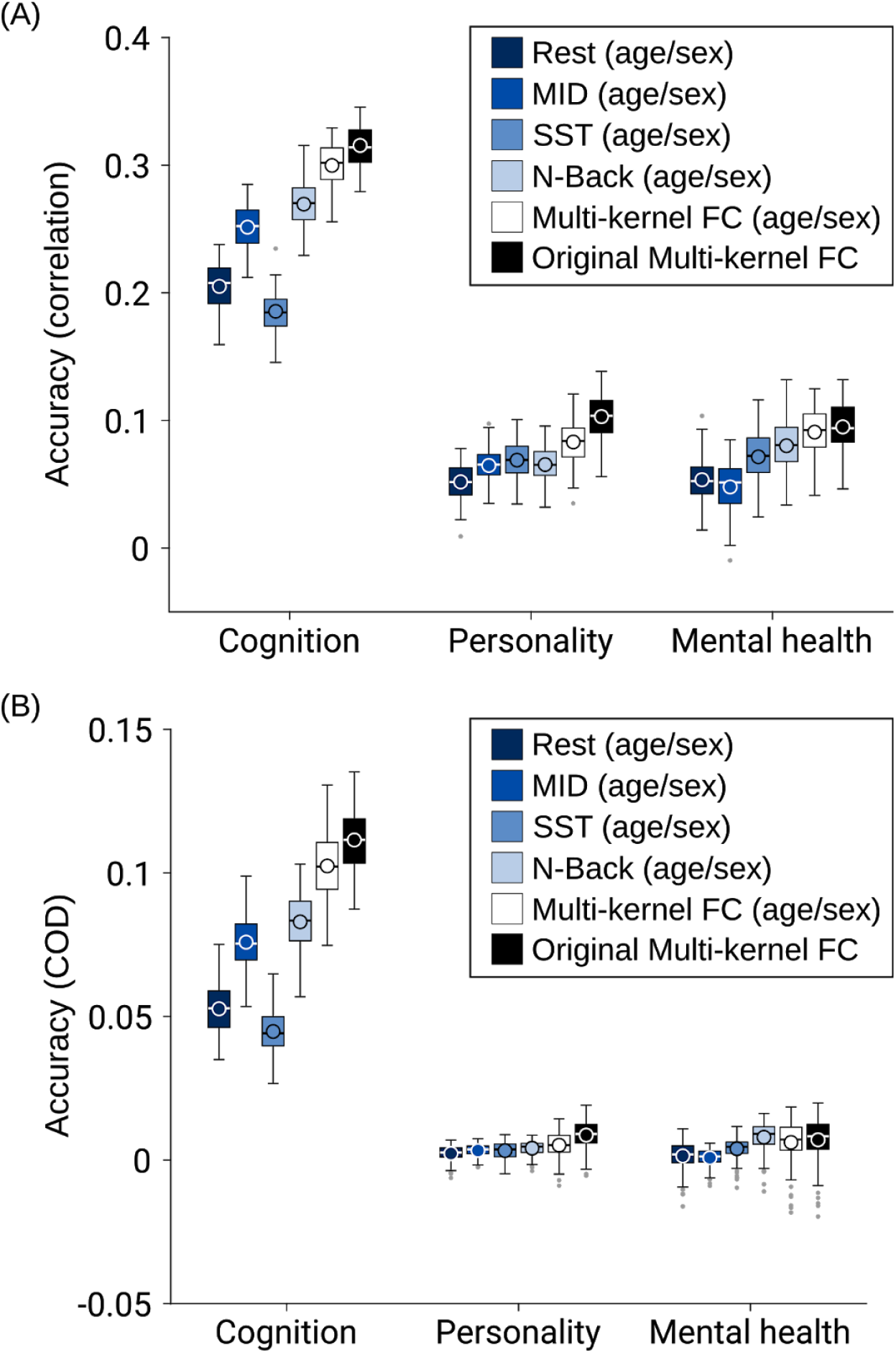
Mean cross-validated prediction performance after regressing out age and sex from the behaviors, compared to the prediction performance of the original multi-kernel FC regression model (as shown in main text) without the regression of age and sex. (A) Accuracy as measured by Pearson’s correlation between observed and predicted values. (B) Accuracy as measured by the coefficient of determination (COD).

**Figures S17.**
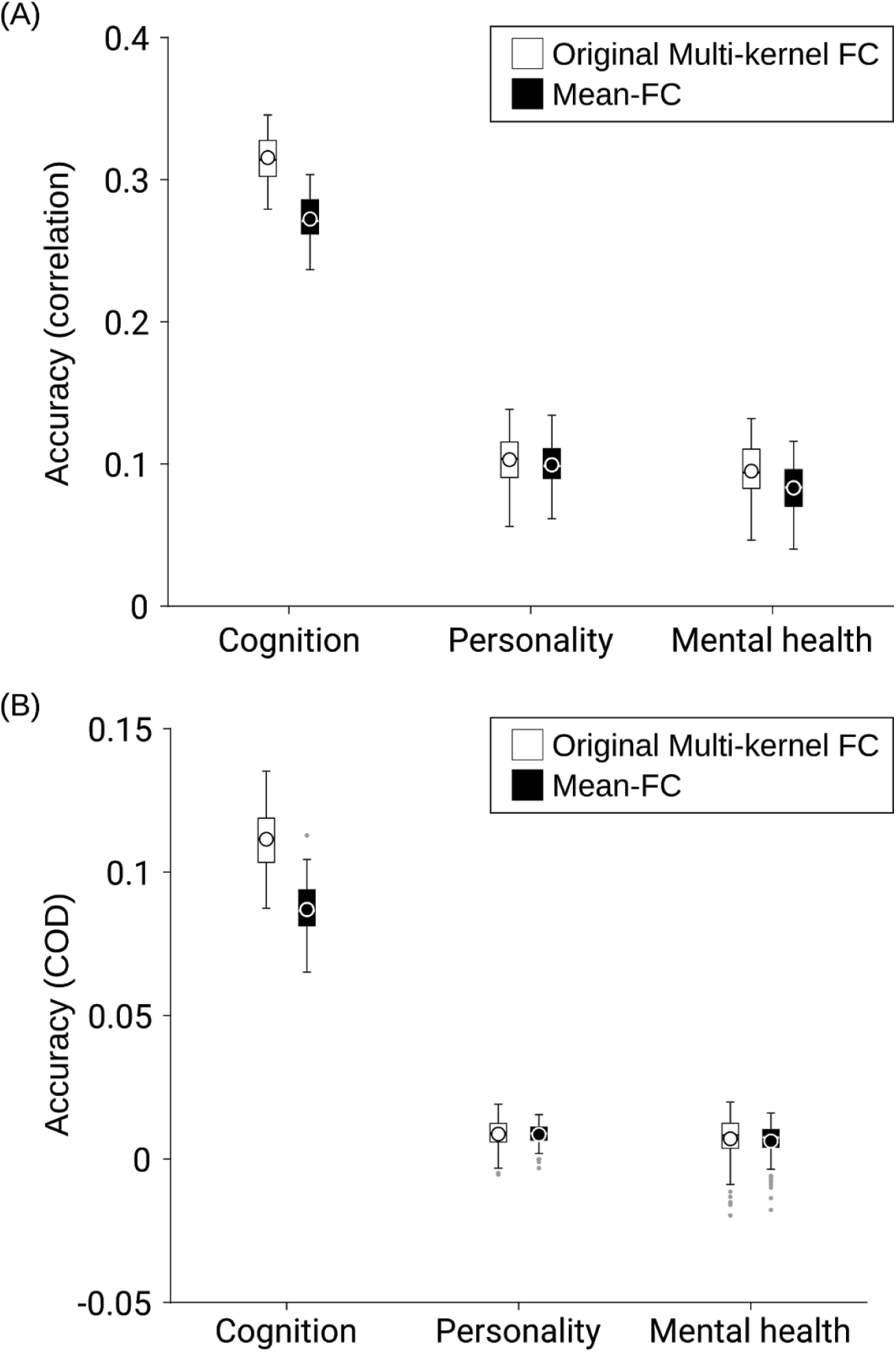
Mean cross-validated prediction performance obtained by the original multi-kernel FC regression model (as shown in main text) and kernel ridge regression using FC averaged across all four brain states (mean-FC). (A) Accuracy as measured by Pearson's correlations between observed and predicted values. (B) Accuracy as measured by the coefficient of determination (COD).

**Figure S18.**
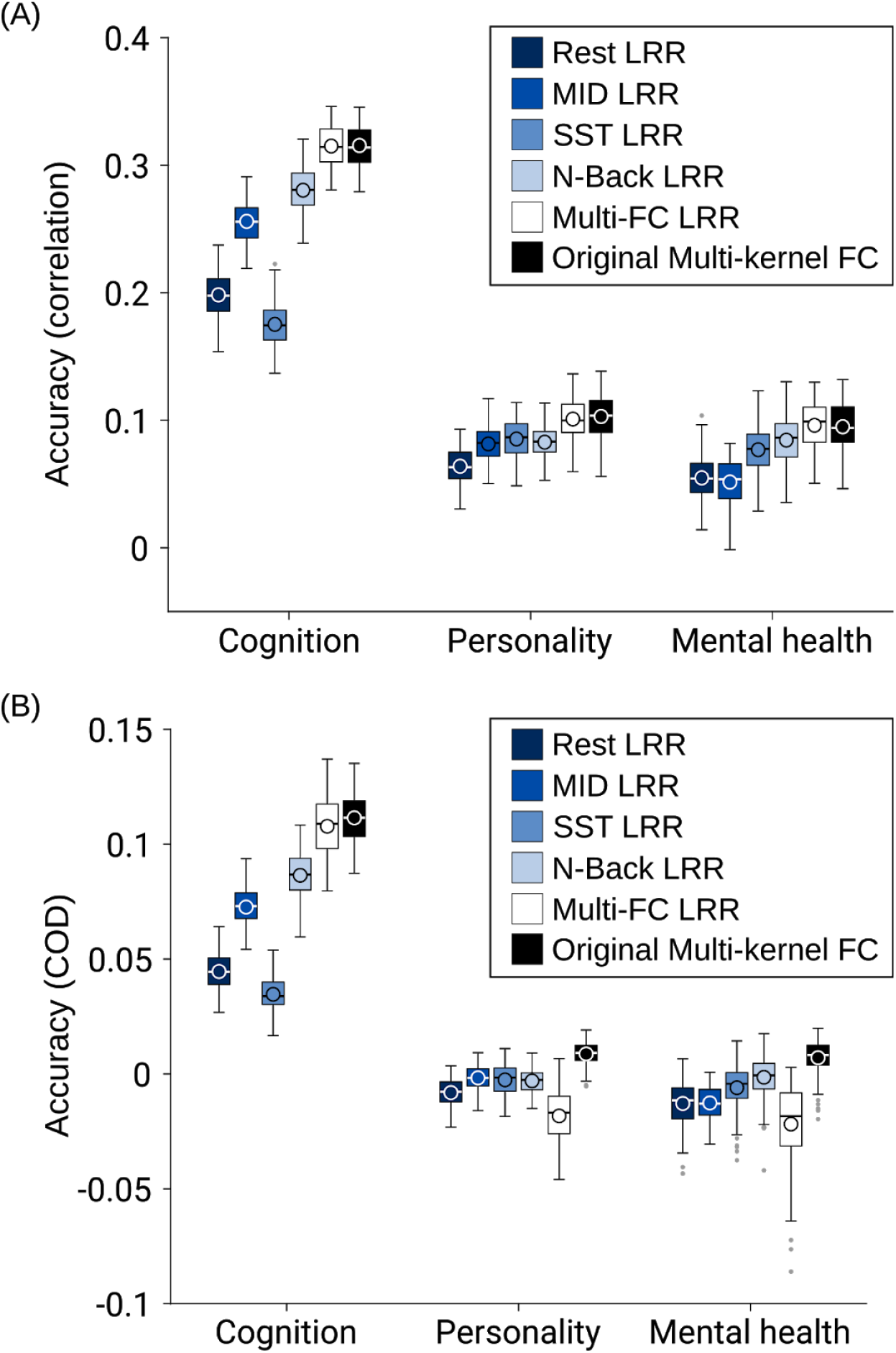
Mean cross-validated prediction performance using linear ridge regression (LRR) and the original multi-kernel FC regression model (as shown in main text). (A) Accuracy as measured by Pearson’s correlations between observed and predicted values. (B) Accuracy as measured by the coefficient of determination (COD).

## References

Achenbach, T. and Rescorla, L., 2013. Achenbach System of Empirically Based Assessment. *In*: F.R. Volkmar, ed. Encyclopedia of Autism Spectrum Disorders. New York, NY: Springer New York, 31–39.

Acker, W. and Acker, C., 1982. Bexley Maudsley automated processing screening and Bexley Maudsley category sorting test manual. Windsor, England: NFER-Nelson.

Baker, J.T., Dillon, D.G., Patrick, L.M., Roffman, J.L., Brady, R.O., Jr, Pizzagalli, D.A., Öngür, D., and Holmes, A.J., 2019. Functional connectomics of affective and psychotic pathology. Proceedings of the National Academy of Sciences of the United States of America, 116 (18), 9050–9059.

Baker, J.T., Holmes, A.J., Masters, G.A., Yeo, B.T.T., Krienen, F., Buckner, R.L., and Öngür, D., 2014. Disruption of cortical association networks in schizophrenia and psychotic bipolar disorder. JAMA psychiatry, 71 (2), 109–118.

Balodis, I.M., Kober, H., Worhunsky, P.D., Stevens, M.C., Pearlson, G.D., and Potenza, M.N., 2012. Diminished frontostriatal activity during processing of monetary rewards and losses in pathological gambling. Biological psychiatry, 71 (8), 749–757.

Barch, D.M., Albaugh, M.D., Avenevoli, S., Chang, L., Clark, D.B., Glantz, M.D., Hudziak, J.J., Jernigan, T.L., Tapert, S.F., Yurgelun-Todd, D., Alia-Klein, N., Potter, A.S., Paulus, M.P., Prouty, D., Zucker, R.A., and Sher, K.J., 2018. Demographic, physical and mental health assessments in the adolescent brain and cognitive development study: Rationale and description. Developmental cognitive neuroscience, 32, 55–66.

Beck, A., Schlagenhauf, F., Wüstenberg, T., Hein, J., Kienast, T., Kahnt, T., Schmack, K., Hägele, C., Knutson, B., Heinz, A., and Wrase, J., 2009. Ventral striatal activation during reward anticipation correlates with impulsivity in alcoholics. Biological psychiatry, 66 (8), 734–742.

Bertolero, M.A., Yeo, B.T.T., Bassett, D.S., and D’Esposito, M., 2018. A mechanistic model of connector hubs, modularity and cognition. Nature human behaviour, 2 (10), 765–777.

Binder, J.R., Frost, J.A., Hammeke, T.A., Cox, R.W., Rao, S.M., and Prieto, T., 1997. Human brain language areas identified by functional magnetic resonance imaging. The Journal of neuroscience: the official journal of the Society for Neuroscience, 17 (1), 353–362.

Bjork, J.M., Knutson, B., Fong, G.W., Caggiano, D.M., Bennett, S.M., and Hommer, D.W., 2004. Incentive-elicited brain activation in adolescents: similarities and differences from young adults. The Journal of neuroscience: the official journal of the Society for Neuroscience, 24 (8), 1793–1802.

Bouckaert, R.R. and Frank, E., 2004. Evaluating the Replicability of Significance Tests for Comparing Learning Algorithms. *In*: Advances in Knowledge Discovery and Data Mining. Springer Berlin Heidelberg, 3–12.

Braga, R.M., Di Nicola, L.M., and Buckner, R.L., 2019. Situating the Left-Lateralized Language Network in the Broader Organization of Multiple Specialized Large-Scale Distributed Networks. bioRxiv.

Buckholtz, J.W., Treadway, M.T., Cowan, R.L., Woodward, N.D., Li, R., Ansari, M.S., Baldwin, R.M., Schwartzman, A.N., Shelby, E.S., Smith, C.E., Kessler, R.M., and Zald, D.H., 2010. Dopaminergic network differences in human impulsivity. Science, 329 (5991), 532.

Bzdok, D., Eickenberg, M., Grisel, O., Thirion, B., and Varoquaux, G., 2015. Semi-Supervised Factored Logistic Regression for High-Dimensional Neuroimaging Data. *In*: C. Cortes, N.D. Lawrence, D.D. Lee, M. Sugiyama, and R. Garnett, eds. Advances in Neural Information Processing Systems 28. Curran Associates, Inc., 3348–3356.

Bzdok, D. and Ioannidis, J.P.A., 2019. Exploration, Inference, and Prediction in Neuroscience and Biomedicine. Trends in neurosciences, 42 (4), 251–262.

Bzdok, D. and Meyer-Lindenberg, A., 2018. Machine Learning for Precision Psychiatry: Opportunities and Challenges. Biological psychiatry. Cognitive neuroscience and neuroimaging, 3 (3), 223–230.

Bzdok, D., Varoquaux, G., Grisel, O., Eickenberg, M., Poupon, C., and Thirion, B., 2016. Formal Models of the Network Co-occurrence Underlying Mental Operations. PLoS computational biology, 12 (6), e1004994.

Carroll, J.B., 2003. Chapter 1 - The Higher-stratum Structure of Cognitive Abilities: Current Evidence Supports g and About Ten Broad Factors. *In*: H. Nyborg, ed. The Scientific Study of General Intelligence. Oxford: Pergamon, 5–21.

Casey, B.J., Cannonier, T., Conley, M.I., Cohen, A.O., Barch, D.M., Heitzeg, M.M., Soules, M.E., Teslovich, T., Dellarco, D.V., Garavan, H., Orr, C.A., Wager, T.D., Banich, M.T., Speer, N.K., Sutherland, M.T., Riedel, M.C., Dick, A.S., Bjork, J.M., Thomas, K.M., Chaarani, B., Mejia, M.H., Hagler, D.J., Jr, Daniela Cornejo, M., Sicat, C.S., Harms, M.P., Dosenbach, N.U.F., Rosenberg, M., Earl, E., Bartsch, H., Watts, R., Polimeni, J.R., Kuperman, J.M., Fair, D.A., Dale, A.M., and ABCD Imaging Acquisition Workgroup, 2018. The Adolescent Brain Cognitive Development (ABCD) study: Imaging acquisition across 21 sites. Developmental cognitive neuroscience, 32, 43–54.

Casey, B.J., Getz, S., and Galvan, A., 2008. The adolescent brain. Developmental review: DR, 28 (1), 62–77.

Caspi, A., Houts, R.M., Belsky, D.W., Goldman-Mellor, S.J., Harrington, H., Israel, S., Meier, M.H., Ramrakha, S., Shalev, I., Poulton, R., and Moffitt, T.E., 2014. The p Factor: One General Psychopathology Factor in the Structure of Psychiatric Disorders? Clinical psychological science, 2 (2), 119–137.

Cohen, J.R. and D’Esposito, M., 2016. The Segregation and Integration of Distinct Brain Networks and Their Relationship to Cognition. The Journal of neuroscience: the official journal of the Society for Neuroscience, 36 (48), 12083–12094.

Cole, M.W., Bassett, D.S., Power, J.D., Braver, T.S., and Petersen, S.E., 2014. Intrinsic and task-evoked network architectures of the human brain. Neuron, 83 (1), 238–251.

Cole, M.W., Repovš, G., and Anticevic, A., 2014. The frontoparietal control system: a central role in mental health. The Neuroscientist: a review journal bringing neurobiology, neurology and psychiatry, 20 (6), 652–664.

Cole, M.W., Reynolds, J.R., Power, J.D., Repovs, G., Anticevic, A., and Braver, T.S., 2013. Multi-task connectivity reveals flexible hubs for adaptive task control. Nature neuroscience, 16 (9), 1348–1355.

Corkin, S., 2002. What’s new with the amnesic patient H.M.? Nature reviews. Neuroscience, 3 (2), 153–160.

Cubillo, A., Halari, R., Smith, A., Taylor, E., and Rubia, K., 2012. A review of fronto-striatal and fronto-cortical brain abnormalities in children and adults with Attention Deficit Hyperactivity Disorder (ADHD) and new evidence for dysfunction in adults with ADHD during motivation and attention. Cortex; a journal devoted to the study of the nervous system and behavior, 48 (2), 194–215.

Cui, Z., Li, H., Xia, C.H., Larsen, B., Adebimpe, A., Baum, G.L., Cieslak, M., Gur, R.E., Gur, R.C., Moore, T.M., Oathes, D.J., Alexander-Bloch, A.F., Raznahan, A., Roalf, D.R., Shinohara, R.T., Wolf, D.H., Davatzikos, C., Bassett, D.S., Fair, D.A., Fan, Y., and Satterthwaite, T.D., 2020. Individual Variation in Functional Topography of Association Networks in Youth. Neuron.

Dale, A.M., Fischl, B., and Sereno, M.I., 1999. Cortical surface-based analysis. I. Segmentation and surface reconstruction. NeuroImage, 9 (2), 179–194.

Dalley, J.W., Mar, A.C., Economidou, D., and Robbins, T.W., 2008. Neurobehavioral mechanisms of impulsivity: fronto-striatal systems and functional neurochemistry. Pharmacology, biochemistry, and behavior, 90 (2), 250–260.

Dosenbach, N.U.F., Nardos, B., Cohen, A.L., Fair, D.A., Power, J.D., Church, J.A., Nelson, S.M., Wig, G.S., Vogel, A.C., Lessov-Schlaggar, C.N., Barnes, K.A., Dubis, J.W., Feczko, E., Coalson, R.S., Pruett, J.R., Jr, Barch, D.M., Petersen, S.E., and Schlaggar, B.L., 2010. Prediction of individual brain maturity using fMRI. Science, 329 (5997), 1358–1361.

Dubois, J., Galdi, P., Han, Y., Paul, L.K., and Adolphs, R., 2018. Resting-state functional brain connectivity best predicts the personality dimension of openness to experience. Personality neuroscience, 1.

Eickhoff, S.B. and Langner, R., 2019. Neuroimaging-based prediction of mental traits: Road to utopia or Orwell? PLoS biology, 17 (11), e3000497.

Elliott, M.L., Knodt, A.R., Cooke, M., Kim, M.J., Melzer, T.R., Keenan, R., Ireland, D., Ramrakha, S., Poulton, R., Caspi, A., Moffitt, T.E., and Hariri, A.R., 2019. General functional connectivity: Shared features of resting-state and task fMRI drive reliable and heritable individual differences in functional brain networks. NeuroImage, 189, 516–532.

Evans, T.M., Kochalka, J., Ngoon, T.J., Wu, S.S., Qin, S., Battista, C., and Menon, V., 2015. Brain Structural Integrity and Intrinsic Functional Connectivity Forecast 6 Year Longitudinal Growth in Children’s Numerical Abilities. The Journal of neuroscience: the official journal of the Society for Neuroscience, 35 (33), 11743–11750.

Fair, D.A., Cohen, A.L., Power, J.D., Dosenbach, N.U.F., Church, J.A., Miezin, F.M., Schlaggar, B.L., and Petersen, S.E., 2009. Functional brain networks develop from a ‘local to distributed’ organization. PLoS computational biology, 5 (5), e1000381.

Farr, O.M., Hu, S., Zhang, S., and Li, C.-S.R., 2012. Decreased saliency processing as a neural measure of Barratt impulsivity in healthy adults. NeuroImage, 63 (3), 1070–1077.

Fedorenko, E., Duncan, J., and Kanwisher, N., 2012. Language-selective and domain-general regions lie side by side within Broca’s area. Current biology: CB, 22 (21), 2059–2062.

Fineberg, N.A., Potenza, M.N., Chamberlain, S.R., Berlin, H.A., Menzies, L., Bechara, A., Sahakian, B.J., Robbins, T.W., Bullmore, E.T., and Hollander, E., 2010. Probing compulsive and impulsive behaviors, from animal models to endophenotypes: a narrative review. Neuropsychopharmacology: official publication of the American College of Neuropsychopharmacology, 35 (3), 591–604.

Finn, E.S., Shen, X., Scheinost, D., Rosenberg, M.D., Huang, J., Chun, M.M., Papademetris, X., and Constable, R.T., 2015. Functional connectome fingerprinting: identifying individuals using patterns of brain connectivity. Nature neuroscience, 18 (11), 1664–1671.

Fischl, B., Liu, A., and Dale, A.M., 2001. Automated manifold surgery: constructing geometrically accurate and topologically correct models of the human cerebral cortex. IEEE transactions on medical imaging, 20 (1), 70–80.

Fischl, B., Salat, D.H., Busa, E., Albert, M., Dieterich, M., Haselgrove, C., van der Kouwe, A., Killiany, R., Kennedy, D., Klaveness, S., Montillo, A., Makris, N., Rosen, B., and Dale, A.M., 2002. Whole brain segmentation: automated labeling of neuroanatomical structures in the human brain. Neuron, 33 (3), 341–355.

Fischl, B., Sereno, M.I., and Dale, A.M., 1999. II: Inflation, Flattening, and a Surface-Based Coordinate System. NeuroImage, 9, 195–207.

Fischl, B., Sereno, M.I., Tootell, R.B., and Dale, A.M., 1999. High-resolution intersubject averaging and a coordinate system for the cortical surface. Human brain mapping, 8 (4), 272–284.

Fong, A.H.C., Yoo, K., Rosenberg, M.D., Zhang, S., Li, C.-S.R., Scheinost, D., Constable, R.T., and Chun, M.M., 2019. Dynamic functional connectivity during task performance and rest predicts individual differences in attention across studies. NeuroImage, 188, 14–25.

Freiwald, W.A. and Tsao, D.Y., 2010. Functional compartmentalization and viewpoint generalization within the macaque face-processing system. Science, 330 (6005), 845–851.

Galvan, A., Hare, T.A., Parra, C.E., Penn, J., Voss, H., Glover, G., and Casey, B.J., 2006. Earlier development of the accumbens relative to orbitofrontal cortex might underlie risk-taking behavior in adolescents. The Journal of neuroscience: the official journal of the Society for Neuroscience, 26 (25), 6885–6892.

Gao, S., Greene, A.S., Constable, R.T., and Scheinost, D., 2019. Combining multiple connectomes improves predictive modeling of phenotypic measures. NeuroImage, 201, 116038.

Gee, D.G., Humphreys, K.L., Flannery, J., Goff, B., Telzer, E.H., Shapiro, M., Hare, T.A., Bookheimer, S.Y., and Tottenham, N., 2013. A developmental shift from positive to negative connectivity in human amygdala-prefrontal circuitry. The Journal of neuroscience: the official journal of the Society for Neuroscience, 33 (10), 4584–4593.

Golchert, J., Smallwood, J., Jefferies, E., Liem, F., Huntenburg, J.M., Falkiewicz, M., Lauckner, M.E., Oligschläger, S., Villringer, A., and Margulies, D.S., 2017. In need of constraint: Understanding the role of the cingulate cortex in the impulsive mind. NeuroImage, 146, 804–813.

Goodkind, M., Eickhoff, S.B., Oathes, D.J., Jiang, Y., Chang, A., Jones-Hagata, L.B., Ortega, B.N., Zaiko, Y.V., Roach, E.L., Korgaonkar, M.S., Grieve, S.M., Galatzer-Levy, I., Fox, P.T., and Etkin, A., 2015. Identification of a common neurobiological substrate for mental illness. JAMA psychiatry, 72 (4), 305–315.

Gordon, E.M., Laumann, T.O., Adeyemo, B., Huckins, J.F., Kelley, W.M., and Petersen, S.E., 2016. Generation and Evaluation of a Cortical Area Parcellation from Resting-State Correlations. Cerebral cortex, 26 (1), 288–303.

Gratton, C., Laumann, T.O., Nielsen, A.N., Greene, D.J., Gordon, E.M., Gilmore, A.W., Nelson, S.M., Coalson, R.S., Snyder, A.Z., Schlaggar, B.L., Dosenbach, N.U.F., and Petersen, S.E., 2018. Functional Brain Networks Are Dominated by Stable Group and Individual Factors, Not Cognitive or Daily Variation. Neuron, 98 (2), 439–452.e5.

Greene, A.S., Gao, S., Scheinost, D., and Constable, R.T., 2018. Task-induced brain state manipulation improves prediction of individual traits. Nature communications, 9 (1), 2807.

Greve, D.N. and Fischl, B., 2009. Accurate and robust brain image alignment using boundary-based registration. NeuroImage, 48 (1), 63–72.

Hagler, D.J., Jr, Hatton, S., Cornejo, M.D., Makowski, C., Fair, D.A., Dick, A.S., Sutherland, M.T., Casey, B.J., Barch, D.M., Harms, M.P., Watts, R., Bjork, J.M., Garavan, H.P., Hilmer, L., Pung, C.J., Sicat, C.S., Kuperman, J., Bartsch, H., Xue, F., Heitzeg, M.M., Laird, A.R., Trinh, T.T., Gonzalez, R., Tapert, S.F., Riedel, M.C., Squeglia, L.M., Hyde, L.W., Rosenberg, M.D., Earl, E.A., Howlett, K.D., Baker, F.C., Soules, M., Diaz, J., de Leon, O.R., Thompson, W.K., Neale, M.C., Herting, M., Sowell, E.R., Alvarez, R.P., Hawes, S.W., Sanchez, M., Bodurka, J., Breslin, F.J., Morris, A.S., Paulus, M.P., Simmons, W.K., Polimeni, J.R., van der Kouwe, A., Nencka, A.S., Gray, K.M., Pierpaoli, C., Matochik, J.A., Noronha, A., Aklin, W.M., Conway, K., Glantz, M., Hoffman, E., Little, R., Lopez, M., Pariyadath, V., Weiss, S.R., Wolff-Hughes, D.L., DelCarmen-Wiggins, R., Feldstein Ewing, S.W., Miranda-Dominguez, O., Nagel, B.J., Perrone, A.J., Sturgeon, D.T., Goldstone, A., Pfefferbaum, A., Pohl, K.M., Prouty, D., Uban, K., Bookheimer, S.Y., Dapretto, M., Galvan, A., Bagot, K., Giedd, J., Infante, M.A., Jacobus, J., Patrick, K., Shilling, P.D., Desikan, R., Li, Y., Sugrue, L., Banich, M.T., Friedman, N., Hewitt, J.K., Hopfer, C., Sakai, J., Tanabe, J., Cottler, L.B., Nixon, S.J., Chang, L., Cloak, C., Ernst, T., Reeves, G., Kennedy, D.N., Heeringa, S., Peltier, S., Schulenberg, J., Sripada, C., Zucker, R.A., Iacono, W.G., Luciana, M., Calabro, F.J., Clark, D.B., Lewis, D.A., Luna, B., Schirda, C., Brima, T., Foxe, J.J., Freedman, E.G., Mruzek, D.W., Mason, M.J., Huber, R., McGlade, E., Prescot, A., Renshaw, P.F., Yurgelun-Todd, D.A., Allgaier, N.A., Dumas, J.A., Ivanova, M., Potter, A., Florsheim, P., Larson, C., Lisdahl, K., Charness, M.E., Fuemmeler, B., Hettema, J.M., Maes, H.H., Steinberg, J., Anokhin, A.P., Glaser, P., Heath, A.C., Madden, P.A., Baskin-Sommers, A., Constable, R.T., Grant, S.J., Dowling, G.J., Brown, S.A., Jernigan, T.L., and Dale, A.M., 2019. Image processing and analysis methods for the Adolescent Brain Cognitive Development Study. NeuroImage, 202, 116091.

Haufe, S., Meinecke, F., Görgen, K., Dähne, S., Haynes, J.-D., Blankertz, B., and Bießmann, F., 2014. On the interpretation of weight vectors of linear models in multivariate neuroimaging. NeuroImage, 87, 96–110.

He, T., Kong, R., Holmes, A.J., Nguyen, M., Sabuncu, M.R., Eickhoff, S.B., Bzdok, D., Feng, J., and Yeo, B.T.T., 2020. Deep neural networks and kernel regression achieve comparable accuracies for functional connectivity prediction of behavior and demographics. NeuroImage, 206, 116276.

van den Heuvel, M.P. and Sporns, O., 2011. Rich-club organization of the human connectome. The Journal of neuroscience: the official journal of the Society for Neuroscience, 31 (44), 15775–15786.

Hodes, R.J., Insel, T.R., Landis, S.C., and NIH Blueprint for Neuroscience Research, 2013. The NIH toolbox: setting a standard for biomedical research. Neurology, 80 (11 Suppl 3), S1.

Holmes, A.J. and Patrick, L.M., 2018. The Myth of Optimality in Clinical Neuroscience. Trends in cognitive sciences, 22 (3), 241–257.

Hsu, W.-T., Rosenberg, M.D., Scheinost, D., Constable, R.T., and Chun, M.M., 2018. Resting-state functional connectivity predicts neuroticism and extraversion in novel individuals. Social cognitive and affective neuroscience, 13 (2), 224–232.

Inuggi, A., Sanz-Arigita, E., González-Salinas, C., Valero-García, A.V., García-Santos, J.M., and Fuentes, L.J., 2014. Brain functional connectivity changes in children that differ in impulsivity temperamental trait. Frontiers in behavioral neuroscience, 8, 156.

Jalbrzikowski, M., Larsen, B., Hallquist, M.N., Foran, W., Calabro, F., and Luna, B., 2017. Development of White Matter Microstructure and Intrinsic Functional Connectivity Between the Amygdala and Ventromedial Prefrontal Cortex: Associations With Anxiety and Depression. Biological psychiatry, 82 (7), 511–521.

Jenkinson, M., Bannister, P., Brady, M., and Smith, S., 2002. Improved optimization for the robust and accurate linear registration and motion correction of brain images. NeuroImage, 17 (2), 825–841.

Jentsch, J.D. and Taylor, J.R., 1999. Impulsivity resulting from frontostriatal dysfunction in drug abuse: implications for the control of behavior by reward-related stimuli. Psychopharmacology, 146 (4), 373–390.

Jiang, R., Zuo, N., Ford, J.M., Qi, S., Zhi, D., Zhuo, C., Xu, Y., Fu, Z., Bustillo, J., Turner, J.A., Calhoun, V.D., and Sui, J., 2019. Task-induced brain connectivity promotes the detection of individual differences in brain-behavior relationships. NeuroImage, 116370.

Karcher, N.R., O’Brien, K.J., Kandala, S., and Barch, D.M., 2019. Resting-State Functional Connectivity and Psychotic-like Experiences in Childhood: Results From the Adolescent Brain Cognitive Development Study. Biological psychiatry.

Kebets, V., Holmes, A.J., Orban, C., Tang, S., Li, J., Sun, N., Kong, R., Poldrack, R.A., and Yeo, B.T.T., 2019. Somatosensory-Motor Dysconnectivity Spans Multiple Transdiagnostic Dimensions of Psychopathology. Biological psychiatry, 86 (10), 779–791.

Kessler, R.C., Ormel, J., Petukhova, M., McLaughlin, K.A., Green, J.G., Russo, L.J., Stein, D.J., Zaslavsky, A.M., Aguilar-Gaxiola, S., Alonso, J., Andrade, L., Benjet, C., de Girolamo, G., de Graaf, R., Demyttenaere, K., Fayyad, J., Haro, J.M., Hu, C. yi, Karam, A., Lee, S., Lepine, J.-P., Matchsinger, H., Mihaescu-Pintia, C., Posada-Villa, J., Sagar, R., and Ustün, T.B., 2011. Development of lifetime comorbidity in the World Health Organization world mental health surveys. Archives of general psychiatry, 68 (1), 90–100.

Kong, R., Li, J., Orban, C., Sabuncu, M.R., Liu, H., Schaefer, A., Sun, N., Zuo, X.-N., Holmes, A.J., Eickhoff, S.B., and Yeo, B.T.T., 2019. Spatial Topography of Individual-Specific Cortical Networks Predicts Human Cognition, Personality, and Emotion. Cerebral cortex, 29 (6), 2533–2551.

Kotov, R., Krueger, R.F., Watson, D., Achenbach, T.M., Althoff, R.R., Bagby, R.M., Brown, T.A., Carpenter, W.T., Caspi, A., Clark, L.A., Eaton, N.R., Forbes, M.K., Forbush, K.T., Goldberg, D., Hasin, D., Hyman, S.E., Ivanova, M.Y., Lynam, D.R., Markon, K., Miller, J.D., Moffitt, T.E., Morey, L.C., Mullins-Sweatt, S.N., Ormel, J., Patrick, C.J., Regier, D.A., Rescorla, L., Ruggero, C.J., Samuel, D.B., Sellbom, M., Simms, L.J., Skodol, A.E., Slade, T., South, S.C., Tackett, J.L., Waldman, I.D., Waszczuk, M.A., Widiger, T.A., Wright, A.G.C., and Zimmerman, M., 2017. The Hierarchical Taxonomy of Psychopathology (HiTOP): A dimensional alternative to traditional nosologies. Journal of abnormal psychology, 126 (4), 454–477.

Kozak, M.J. and Cuthbert, B.N., 2016. The NIMH Research Domain Criteria Initiative: Background, Issues, and Pragmatics. Psychophysiology, 53 (3), 286–297.

Krienen, F.M., Yeo, B.T.T., and Buckner, R.L., 2014. Reconfigurable task-dependent functional coupling modes cluster around a core functional architecture. Philosophical transactions of the Royal Society of London. Series B, Biological sciences, 369 (1653), 20130526–20130526.

Laird, A.R., Fox, P.M., Eickhoff, S.B., Turner, J.A., Ray, K.L., McKay, D.R., Glahn, D.C., Beckmann, C.F., Smith, S.M., and Fox, P.T., 2011. Behavioral interpretations of intrinsic connectivity networks. Journal of cognitive neuroscience, 23 (12), 4022–4037.

Lake, E.M.R., Finn, E.S., Noble, S.M., Vanderwal, T., Shen, X., Rosenberg, M.D., Spann, M.N., Chun, M.M., Scheinost, D., and Constable, R.T., 2019. The functional brain organization of an individual allows prediction of measures of social abilities trans-diagnostically in autism and attention/deficit and hyperactivity disorder. Biological psychiatry.

Larsen, B. and Luna, B., 2018. Adolescence as a neurobiological critical period for the development of higher-order cognition. Neuroscience and biobehavioral reviews, 94, 179–195.

Leshem, R. and Glicksohn, J., 2007. The construct of impulsivity revisited. Personality and individual differences, 43 (4), 681–691.

Liégeois, R., Li, J., Kong, R., Orban, C., Van De Ville, D., Ge, T., Sabuncu, M.R., and Yeo, B.T.T., 2019. Resting brain dynamics at different timescales capture distinct aspects of human behavior. Nature communications, 10 (1), 2317.

Li, J., Kong, R., Liégeois, R., Orban, C., Tan, Y., Sun, N., Holmes, A.J., Sabuncu, M.R., Ge, T., and Yeo, B.T.T., 2019. Global signal regression strengthens association between resting-state functional connectivity and behavior. NeuroImage, 196, 126–141.

Loewy, R.L., Therman, S., Manninen, M., Huttunen, M.O., and Cannon, T.D., 2012. Prodromal psychosis screening in adolescent psychiatry clinics. Early intervention in psychiatry, 6 (1), 69–75.

Luciana, M., Bjork, J.M., Nagel, B.J., Barch, D.M., Gonzalez, R., Nixon, S.J., and Banich, M.T., 2018. Adolescent neurocognitive development and impacts of substance use: Overview of the adolescent brain cognitive development (ABCD) baseline neurocognition battery. Developmental cognitive neuroscience, 32, 67–79.

Lynam, D.R., 2013. Development of a short form of the UPPS-P Impulsive Behavior Scale. Unpublished Technical Report.

Maglanoc, L.A., Kaufmann, T., van der Meer, D., Marquand, A.F., Wolfers, T., Jonassen, R., Hilland, E., Andreassen, O.A., Landrø, N.I., and Westlye, L.T., 2019. Brain connectome mapping of complex human traits and their polygenic architecture using machine learning. Biological psychiatry, 0 (0).

Marek, S., Tervo-Clemmens, B., Nielsen, A.N., Wheelock, M.D., Miller, R.L., Laumann, T.O., Earl, E., Foran, W.W., Cordova, M., Doyle, O., Perrone, A., Miranda-Dominguez, O., Feczko, E., Sturgeon, D., Graham, A., Hermosillo, R., Snider, K., Galassi, A., Nagel, B.J., Ewing, S.W.F., Eggebrecht, A.T., Garavan, H., Dale, A.M., Greene, D.J., Barch, D.M., Fair, D.A., Luna, B., and Dosenbach, N.U.F., 2019. Identifying Reproducible Individual Differences in Childhood Functional Brain Networks: An ABCD Study. Developmental cognitive neuroscience, 100706.

Menon, V., 2011. Large-scale brain networks and psychopathology: a unifying triple network model. Trends in cognitive sciences, 15 (10), 483–506.

Menon, V. and Uddin, L.Q., 2010. Saliency, switching, attention and control: a network model of insula function. Brain structure & function, 214 (5-6), 655–667.

Milham, M.P., Craddock, R.C., and Klein, A., 2017. Clinically useful brain imaging for neuropsychiatry: How can we get there? Depression and anxiety, 34 (7), 578–587.

Nadeau, C. and Bengio, Y., 2003. Inference for the Generalization Error. Machine learning, 52 (3), 239–281.

Nomura, E.M., Gratton, C., Visser, R.M., Kayser, A., Perez, F., and D’Esposito, M., 2010. Double dissociation of two cognitive control networks in patients with focal brain lesions. Proceedings of the National Academy of Sciences of the United States of America, 107 (26), 12017–12022.

Nostro, A.D., Müller, V.I., Varikuti, D.P., Pläschke, R.N., Hoffstaedter, F., Langner, R., Patil, K.R., and Eickhoff, S.B., 2018. Predicting personality from network-based resting-state functional connectivity. Brain structure & function, 223 (6), 2699–2719.

Orban, C., Kong, R., Li, J., Chee, M.W.L., and Yeo, B.T.T., 2020. Time of day is associated with paradoxical reductions in global signal fluctuation and functional connectivity. PLoS biology, 18 (2), e3000602.

Pagliaccio, D., Luking, K.R., Anokhin, A.P., Gotlib, I.H., Hayden, E.P., Olino, T.M., Peng, C.-Z., Hajcak, G., and Barch, D.M., 2016. Revising the BIS/BAS Scale to study development: Measurement invariance and normative effects of age and sex from childhood through adulthood. Psychological assessment, 28 (4), 429–442.

Paus, T., Keshavan, M., and Giedd, J.N., 2008. Why do many psychiatric disorders emerge during adolescence? Nature reviews. Neuroscience, 9 (12), 947–957.

Petersen, S.E., Fox, P.T., Posner, M.I., Mintun, M., and Raichle, M.E., 1988. Positron emission tomographic studies of the cortical anatomy of single-word processing. Nature, 331 (6157), 585–589.

Pornpattananangkul, N., Leibenluft, E., Pine, D.S., and Stringaris, A., 2019. Association of Brain Functions in Children With Anhedonia Mapped Onto Brain Imaging Measures. JAMA psychiatry.

Power, J.D., Barnes, K.A., Snyder, A.Z., Schlaggar, B.L., and Petersen, S.E., 2012. Spurious but systematic correlations in functional connectivity MRI networks arise from subject motion. NeuroImage, 59 (3), 2142–2154.

Power, J.D., Fair, D.A., Schlaggar, B.L., and Petersen, S.E., 2010. The development of human functional brain networks. Neuron, 67 (5), 735–748.

Power, J.D., Mitra, A., Laumann, T.O., Snyder, A.Z., Schlaggar, B.L., and Petersen, S.E., 2014. Methods to detect, characterize, and remove motion artifact in resting state fMRI. NeuroImage, 84, 320–341.

Rosenberg, M.D., Finn, E.S., Scheinost, D., Papademetris, X., Shen, X., Constable, R.T., and Chun, M.M., 2016. A neuromarker of sustained attention from whole-brain functional connectivity. Nature neuroscience, 19 (1), 165–171.

Russo, M., Levine, S.Z., Demjaha, A., Di Forti, M., Bonaccorso, S., Fearon, P., Dazzan, P., Pariante, C.M., David, A.S., Morgan, C., Murray, R.M., and Reichenberg, A., 2014. Association between symptom dimensions and categorical diagnoses of psychosis: a cross-sectional and longitudinal investigation. Schizophrenia bulletin, 40 (1), 111–119.

Salehi, M., Karbasi, A., Barron, D.S., Scheinost, D., and Constable, R.T., 2019. Individualized functional networks reconfigure with cognitive state. NeuroImage, 116233.

Satterthwaite, T.D., Ruparel, K., Loughead, J., Elliott, M.A., Gerraty, R.T., Calkins, M.E., Hakonarson, H., Gur, R.C., Gur, R.E., and Wolf, D.H., 2012. Being right is its own reward: load and performance related ventral striatum activation to correct responses during a working memory task in youth. NeuroImage, 61 (3), 723–729.

Satterthwaite, T.D., Vandekar, S.N., Wolf, D.H., Bassett, D.S., Ruparel, K., Shehzad, Z., Craddock, R.C., Shinohara, R.T., Moore, T.M., Gennatas, E.D., Jackson, C., Roalf, D.R., Milham, M.P., Calkins, M.E., Hakonarson, H., Gur, R.C., and Gur, R.E., 2015. Connectome-wide network analysis of youth with Psychosis-Spectrum symptoms. Molecular psychiatry, 20 (12), 1508–1515.

Schaefer, A., Kong, R., Gordon, E.M., Laumann, T.O., Zuo, X.-N., Holmes, A.J., Eickhoff, S.B., and Yeo, B.T.T., 2018. Local-Global Parcellation of the Human Cerebral Cortex from Intrinsic Functional Connectivity MRI. Cerebral cortex, 28 (9), 3095–3114.

Schultz, D.H. and Cole, M.W., 2016. Higher Intelligence Is Associated with Less Task-Related Brain Network Reconfiguration. The Journal of neuroscience: the official journal of the Society for Neuroscience, 36 (33), 8551–8561.

Scoville, W.B. and Milner, B., 1957. Loss of recent memory after bilateral hippocampal lesions. Journal of neurology, neurosurgery, and psychiatry, 20 (1), 11–21.

Ségonne, F., Dale, A.M., Busa, E., Glessner, M., Salat, D., Hahn, H.K., and Fischl, B., 2004. A hybrid approach to the skull stripping problem in MRI. NeuroImage, 22 (3), 1060–1075.

Ségonne, F., Pacheco, J., and Fischl, B., 2007. Geometrically accurate topology-correction of cortical surfaces using nonseparating loops. IEEE transactions on medical imaging, 26 (4), 518–529.

Shannon, B.J., Raichle, M.E., Snyder, A.Z., Fair, D.A., Mills, K.L., Zhang, D., Bache, K., Calhoun, V.D., Nigg, J.T., Nagel, B.J., Stevens, A.A., and Kiehl, K.A., 2011. Premotor functional connectivity predicts impulsivity in juvenile offenders. Proceedings of the National Academy of Sciences of the United States of America, 108 (27), 11241–11245.

Sha, Z., Wager, T.D., Mechelli, A., and He, Y., 2019. Common Dysfunction of Large-Scale Neurocognitive Networks Across Psychiatric Disorders. Biological psychiatry, 85 (5), 379–388.

Shine, J.M., Bissett, P.G., Bell, P.T., Koyejo, O., Balsters, J.H., Gorgolewski, K.J., Moodie, C.A., and Poldrack, R.A., 2016. The Dynamics of Functional Brain Networks: Integrated Network States during Cognitive Task Performance. Neuron, 92 (2), 544–554.

Silvers, J.A., Insel, C., Powers, A., Franz, P., Helion, C., Martin, R.E., Weber, J., Mischel, W., Casey, B.J., and Ochsner, K.N., 2017. vlPFC-vmPFC-Amygdala Interactions Underlie Age-Related Differences in Cognitive Regulation of Emotion. Cerebral cortex, 27 (7), 3502–3514.

Smith, S.M., Fox, P.T., Miller, K.L., Glahn, D.C., Fox, P.M., Mackay, C.E., Filippini, N., Watkins, K.E., Toro, R., Laird, A.R., and Beckmann, C.F., 2009. Correspondence of the brain’s functional architecture during activation and rest. Proceedings of the National Academy of Sciences of the United States of America, 106 (31), 13040–13045.

Smith, S.M., Jenkinson, M., Woolrich, M.W., Beckmann, C.F., Behrens, T.E.J., Johansen-Berg, H., Bannister, P.R., De Luca, M., Drobnjak, I., Flitney, D.E., Niazy, R.K., Saunders, J., Vickers, J., Zhang, Y., De Stefano, N., Brady, J.M., and Matthews, P.M., 2004. Advances in functional and structural MR image analysis and implementation as FSL. NeuroImage, 23 Suppl 1, S208–19.

Spear, L.P., 2013. Adolescent neurodevelopment. The Journal of adolescent health: official publication of the Society for Adolescent Medicine, 52 (2 Suppl 2), S7–13.

Sripada, C., Rutherford, S., Angstadt, M., Thompson, W.K., Luciana, M., Weigard, A., Hyde, L.H., and Heitzeg, M., 2019. Prediction of neurocognition in youth from resting state fMRI. Molecular psychiatry.

Steinberg, L., 2005. Cognitive and affective development in adolescence. Trends in cognitive sciences, 9 (2), 69–74.

Strauss, E., Sherman, E.M.S., and Spreen, O., 2006. A Compendium of Neuropsychological Tests: Administration, Norms, and Commentary. Oxford University Press.

Supekar, K., Musen, M., and Menon, V., 2009. Development of large-scale functional brain networks in children. PLoS biology, 7 (7), e1000157.

Swartz, J.R., Carrasco, M., Wiggins, J.L., Thomason, M.E., and Monk, C.S., 2014. Age-related changes in the structure and function of prefrontal cortex-amygdala circuitry in children and adolescents: a multi-modal imaging approach. NeuroImage, 86, 212–220.

Tamminga, C.A., Ivleva, E.I., Keshavan, M.S., Pearlson, G.D., Clementz, B.A., Witte, B., Morris, D.W., Bishop, J., Thaker, G.K., and Sweeney, J.A., 2013. Clinical phenotypes of psychosis in the Bipolar-Schizophrenia Network on Intermediate Phenotypes (B-SNIP). The American journal of psychiatry, 170 (11), 1263–1274.

Uddin, L.Q., Supekar, K., Lynch, C.J., Khouzam, A., Phillips, J., Feinstein, C., Ryali, S., and Menon, V., 2013. Salience network-based classification and prediction of symptom severity in children with autism. JAMA psychiatry, 70 (8), 869–879.

Van Leijenhorst, L., Gunther Moor, B., Op de Macks, Z.A., Rombouts, S.A.R.B., Westenberg, P.M., and Crone, E.A., 2010. Adolescent risky decision-making: neurocognitive development of reward and control regions. NeuroImage, 51 (1), 345–355.

Varoquaux, G., Raamana, P.R., Engemann, D.A., Hoyos-Idrobo, A., Schwartz, Y., and Thirion, B., 2017. Assessing and tuning brain decoders: Cross-validation, caveats, and guidelines. NeuroImage, 145 (Pt B), 166–179.

Virtanen, P., Gommers, R., Oliphant, T.E., Haberland, M., Reddy, T., Cournapeau, D., Burovski, E., Peterson, P., Weckesser, W., Bright, J., van der Walt, S.J., Brett, M., Wilson, J., Millman, K.J., Mayorov, N., Nelson, A.R.J., Jones, E., Kern, R., Larson, E., Carey, C.J., Polat, İ., Feng, Y., Moore, E.W., VanderPlas, J., Laxalde, D., Perktold, J., Cimrman, R., Henriksen, I., Quintero, E.A., Harris, C.R., Archibald, A.M., Ribeiro, A.H., Pedregosa, F., van Mulbregt, P., and SciPy 1.0 Contributors, 2020. SciPy 1.0: fundamental algorithms for scientific computing in Python. Nature methods.

Volkow, N.D., Koob, G.F., Croyle, R.T., Bianchi, D.W., Gordon, J.A., Koroshetz, W.J., Pérez-Stable, E.J., Riley, W.T., Bloch, M.H., Conway, K., Deeds, B.G., Dowling, G.J., Grant, S., Howlett, K.D., Matochik, J.A., Morgan, G.D., Murray, M.M., Noronha, A., Spong, C.Y., Wargo, E.M., Warren, K.R., and Weiss, S.R.B., 2018. The conception of the ABCD study: From substance use to a broad NIH collaboration. Developmental cognitive neuroscience, 32, 4–7.

Wang, D., Li, M., Wang, M., Schoeppe, F., Ren, J., Chen, H., Öngür, D., Baker, J.T., and Liu, H., 2018. Individual-specific functional connectivity markers track dimensional and categorical features of psychotic illness. Molecular psychiatry.

Warren, D.E., Power, J.D., Bruss, J., Denburg, N.L., Waldron, E.J., Sun, H., Petersen, S.E., and Tranel, D., 2014. Network measures predict neuropsychological outcome after brain injury. Proceedings of the National Academy of Sciences of the United States of America, 111 (39), 14247–14252.

Wechsler, D., 2014. Wechsler intelligence scale for children--Fifth Edition (WISC-V). Bloomington, MN: Pearson.

Whitfield-Gabrieli, S. and Ford, J.M., 2012. Default mode network activity and connectivity in psychopathology. Annual review of clinical psychology, 8, 49–76.

Xia, C.H., Ma, Z., Ciric, R., Gu, S., Betzel, R.F., Kaczkurkin, A.N., Calkins, M.E., Cook, P.A., García de la Garza, A., Vandekar, S.N., Cui, Z., Moore, T.M., Roalf, D.R., Ruparel, K., Wolf, D.H., Davatzikos, C., Gur, R.C., Gur, R.E., Shinohara, R.T., Bassett, D.S., and Satterthwaite, T.D., 2018. Linked dimensions of psychopathology and connectivity in functional brain networks. Nature communications, 9 (1), 3003.

Yeo, B.T.T., Krienen, F.M., Eickhoff, S.B., Yaakub, S.N., Fox, P.T., Buckner, R.L., Asplund, C.L., and Chee, M.W.L., 2015. Functional Specialization and Flexibility in Human Association Cortex. Cerebral cortex, 25 (10), 3654–3672.

Yeo, B.T.T., Krienen, F.M., Sepulcre, J., Sabuncu, M.R., Lashkari, D., Hollinshead, M., Roffman, J.L., Smoller, J.W., Zollei, L., Polimeni, J.R., Fischl, B., Liu, H., and Buckner, R.L., 2011. The organization of the human cerebral cortex estimated by intrinsic functional connectivity. Journal of neurophysiology, 106 (3), 1125–1165.

Yoo, K., Rosenberg, M.D., Hsu, W.-T., Zhang, S., Li, C.-S.R., Scheinost, D., Constable, R.T., and Chun, M.M., 2018. Connectome-based predictive modeling of attention: Comparing different functional connectivity features and prediction methods across datasets. NeuroImage, 167, 11–22.

Youngstrom, E.A., Murray, G., Johnson, S.L., and Findling, R.L., 2013. The 7 up 7 down inventory: a 14-item measure of manic and depressive tendencies carved from the General Behavior Inventory. Psychological assessment, 25 (4), 1377–1383.

Zuo, N., Yang, Z., Liu, Y., Li, J., and Jiang, T., 2018. Core networks and their reconfiguration patterns across cognitive loads. Human brain mapping, 39 (9), 3546–3557.

